# Optimized Kakadu Plum Extracts Inhibit Intracellular Oxidative Stress in Canine Small Intestinal Cell Model

**DOI:** 10.64898/2026.08.16.745076

**Authors:** Yidan He, Xuefu Zhou, Antonio Celentano, Nicola Cirillo, Long Cheng, Zhongxiang Fang, Pangzhen Zhang

**Affiliations:** School of Agriculture, Food and Ecosystem Sciences, Faculty of Science, the University of Melbourne, Melbourne, Victoria, Australia; Melbourne Dental School, Faculty of Medicine, Dentistry and Health Sciences, the University of Melbourne, Melbourne, Victoria, Australia; Department of Clinical and Experimental Medicine, University of Foggia, Foggia, Italy; School of Agriculture, Food and Ecosystem Sciences, Faculty of Science, the University of Melbourne, Dookie campus, Victoria, Australia

**Author notes:** Corresponding author: Pangzhen Zhang.

**Keywords:** Kakadu plum extract, Reactive oxygen species (ROS), Intracellular oxidative stress, Network pharmacology, dog food, oxidative stress

## Abstract

Kakadu plum (*Terminalia ferdinandiana*), an Australian native fruit, is among the richest known dietary sources of vitamin C and hydrolysable tannins, yet its capacity to protect the intestinal epithelium against oxidative stress remains largely unexplored. This study optimised the extraction of bioactive compounds from freeze-dried Kakadu plum powder and evaluated their antioxidant activity using both chemical and cellular antioxidant *in vitro* assay. Phenolic compounds were extracted using three solvents (water, 80% ethanol, and 80% methanol) combined with shaking, ultrasound, or microwave assistance. Solvent, rather than processing technique, was the dominant determinant of antioxidant capacity: ethanol and methanol maximised total phenolic content, total flavonoid content, and DPPH radical-scavenging activity, whereas water extracts showed the highest ferric-reducing antioxidant power. Twenty-four phenolic compounds identified by HPLC-ESI-QTOF-MS/MS were mapped by network pharmacology to nine core oxidative-stress targets, and cross-species molecular docking predicted conserved binding of key phenolics to canine orthologs of PTGS2 and MMP2. In an H_2_O_2_-induced oxidative-stress *in vitro* cell model using canine small intestinal epithelial cells, both water (≤25 µg/mL) and ethanol (≤250 µg/mL) extracts significantly suppressed intracellular reactive oxygen species (ROS) in a dose-dependent manner, with the ethanol extract effective across a wider concentration range. This work demonstrated that Kakadu plum extract could be a promising natural, multi-target antioxidant ingredient for canine intestinal health, and provided a reference for future *in vivo* research.

## 1. Introduction

Kakadu plum (*Terminalia ferdinandiana*), a fruit native to northern Australia and traditionally used by Aboriginal communities as both food and medicine, has attracted growing scientific and commercial interest as a functional-food ingredient (Chaliha, 2018). The fruit is distinguished by an exceptionally high ascorbic acid (vitamin C, 120.0-224.9 mg/g dry weight) content and an abundance of hydrolysable tannins, ellagitannins, and phenolic acids, placing it among the most antioxidant-rich fruits characterised to date (Konczak et al., 2014; J. Li et al., 2023). These phytochemicals underpin a broad range of reported bioactivities and have prompted the incorporation of Kakadu plum into beverages, nutraceuticals, cosmetics, and natural preservative systems, and premium pet foods (Bobasa et al., 2022). Despite the potential use of Kakadu plum, its extraction and conversion into bioactive-enriched ingredients is underdeveloped. The recovery of phenolics from plant matrices is strongly influenced by the extraction solvent and the assisting technology, which together govern the release of free and bound phenolics (Hossain et al., 2021). Techniques such as ultrasound- and microwave-assisted extraction have been used to release tannins and other bound phenolics from related matrices (Hernández-Corroto et al., 2022). However, how solvent polarity and processing technique jointly affect the antioxidant profile of Kakadu plum across its free, bound, and total phenolic fractions has not been systematically studied.

Oxidative stress, arising from an imbalance between reactive oxygen species (ROS) production and antioxidant defence, is a central driver of intestinal epithelial injury, barrier dysfunction, and inflammatory bowel disease (IBD) (Sies et al., 2022). Diet-derived polyphenols can counteract this imbalance by scavenging radicals, chelating transition metals, and modulating redox-sensitive signalling. Kakadu plum extracts have shown antioxidant, anti-inflammatory, antimicrobial, anticancer, and hepatoprotective activities across a range of chemical and cellular models (Akter et al., 2021; Alwis et al., 2025; Ramadhania et al., 2022), and their phenolics are extensively biotransformed during colonic fermentation, implying direct interaction with the gut epithelium (Adiamo et al., 2024). Nonetheless, most of the evidence *in vitro* at cellular level derives from cancer or non-intestinal cell lines, and the ability of Kakadu plum phenolics to protect normal intestinal epithelium from oxidative stress remains to be explored. Furthermore, to the best of our knowledge, no HLJOLJ-induced oxidative stress model has been established for canine intestinal epithelial cells.

To address these gaps, the present study aimed to 1. optimise the solvent and processing conditions for extracting phenolics from Kakadu plum and 2. characterise their antioxidant capacity and phenolic profile via HPLC-DAD-ESI-QTOF-MS/MS. The integrated network pharmacology, ADMET, cross-species ortholog-mapping, and molecular-docking workflow were employed to predict the oxidative-stress targets engaged by the identified phenolics and their conservation in the dog. The immortalised canine small intestinal epithelial cell line and HLJOLJ was used to induce cellular oxidative stress. The intracellular antioxidant activity of the optimised Kakadu plum extracts was validated in the HLJOLJ-triggered cell model. This research would provide mechanistic insight in developing Kakadu plum as a potential functional antioxidant ingredient for canine intestinal health.

## 2. Materials and Methods

### 2.1. Materials and equipment

Commercial freeze-dried Kakadu plum fruit powder (13 µm, filtered through a 1000-mesh screen) was purchased from Traditional Homeland Enterprise (Batch Number 16145 from the Northern Territory in Australia, T.H.E., Victoria, Australia). Ethanol, methanol, Folin-Ciocalteu (FCR), gallic acid, sodium carbonate, sodium nitrite, aluminium chloride, sodium hydroxide, catechin, 2,2-diphenyl-1-picrylhydrazyl (DPPH), Trolox, 2,4,6-tripyridyl-s-triazine (TPTZ), Fe [II] sulfate heptahydrate, Hanks’ balanced salt solution (HBSS) without phenol red (H8264), white wall 96-well microplate, dark clear-bottom 96-well microplate, 0.45 µm and 0.22 µm nylon syringe filters were sourced from Sigma-Aldrich (NSW, Australia). T25 polystyrene flasks (Sarstedt AG & Co, Nümbrecht, Germany), epithelial cell medium Kit (#M6621) and 0.05% (w/v) trypsin-EDTA (#6915) were sourced from Cell Biologics Inc., Chicago, USA. ViaLight® Plus Cell Proliferation and Cytotoxicity BioAssay Kit (#LT07, Lonza Bioscience, NSW, Australia), hydrogen peroxide solution 30% w/w were purchased from ChemSupply Australia, SA, Australia).

### 2.2. Extraction of phenolic compounds

To determine the impact of solvent systems and extraction technologies on the physicochemical characteristics of Kakadu plum, the freeze-dried Kakadu plum powder was extracted using different solvents (i.e., ethanol, methanol, and water) assisted by different processing technologies (i.e., microwave, shaking, and ultrasound, independently).

#### 2.2.1. Extraction of free phenolics

The extraction of free phenolics was conducted according to the methods described by Konczak et al. (2014), with slight modifications. The mixture of KP freeze-dried powder and solvents at a ratio of 1:10 (w/v) was vortexed for 3 min and then assisted by different extraction technologies. The extraction solvent included 80% methanol (v/v), 80% ethanol (v/v), deionized water, and the processing technologies included shaking (20 min shaking at room temperature), ultrasound (20 min ultrasound) (Hernández-Corroto et al., 2022), and microwave (600 W microwave for 1 min and shaken for 19 min) (Theocharis & Andlauer, 2013). After the treatment, the mixture was centrifuged (4 °C, 10 min, 5000 g) and the supernatant was collected, freeze-dried, and stored at -20 °C until analysis, while the pellets were retained for bound phenolics extraction.

#### 2.2.2. Extraction of bound phenolics

The extraction of bound phenolic compounds was conducted according to the methods described by Leonard et al. (2021) with slight alterations, using the pellets obtained above. The pellets were mixed with 2 M HCl at a ratio of 1:10 (w/v, pellets: HCl) and incubated at 99 °C for one hour. An equal volume of ethyl acetate was then added and thoroughly mixed. After phase separation, the ethyl acetate fractions were collected and rotary evaporated to dryness at 40 °C. The dried extracts were reconstituted with methanol, freeze-dried, and stored at -20 °C until analysis.

### 2.3. Physicochemical characteristics of the extract of Kakadu plum

#### 2.3.1. Total phenolic content (TPC) assay

The TPC of extracts was evaluated using the Folin-Ciocalteu reagent (FCR) assay according to (Amiri et al., 2023) with minor modification. The freeze-dried extract was reconstituted with methanol and filtered with 0.45µm nylon syringe filters. The extracts were mixed with 0.2 N Folin-Ciocalteu (FCR) and saturated sodium carbonate solution (7.5% w/v), followed with 30 min incubation in the dark at room temperature. The absorbance at 765 nm was measured using the Varioskan LUX Multimode Microplate Reader (Thermo Fisher Scientific Australia Pty Ltd, VIC, Australia). The TPC was expressed as mg gallic acid equivalents (GAE) per g dry weight of the freeze-dried extract (mg GAE/g DW), based on a gallic acid standard curve against the blank control (methanol).

#### 2.3.2. Total flavonoid content (TFC) assay

The TFC of the extract was measured using the aluminium chloride colourimetric method according to Amiri et al. (2023) with a slight modification. The above reconstituted methanol extracts and 5% (w/v) of sodium nitrite solution was incubated for 5 min at room temperature before the addition of 10% (w/v) of aluminium chloride solution and incubation for 6 min. Then, the 0.5 M sodium hydroxide solution was added, and the mixture was incubated in the dark for 30 min at room temperature. The absorbance at 510 nm was measured using the Varioskan LUX Multimode Microplate Reader. The TFC was expressed as mg of catechin equivalent (CAE) per g dry weight of the freeze-dried extract (mg CAE/g DW), based on the catechin standard curve against a blank control.

#### 2.3.3. DPPH radical scavenging capacity assay

Determination of DPPH radical scavenging capacity was performed according to Akter et al. (2019) with slight alterations. The above reconstituted methanol extracts and DPPH solution was incubated in the dark at room temperature for one hour. The absorbance at 517 nm was measured using the Varioskan LUX Multimode Microplate Reader. The DPPH radical scavenging capacity was expressed as mg Trolox equivalent per g dry weight of the freeze-dried extract (mg TE/g DW), based on the Trolox standard curve against a blank control.

#### 2.3.4. Ferric reducing antioxidant power (FRAP) assay

The FRAP assay was performed according to the method described by Konczak et al. (2010) with minor modifications. The FRAP reagent was freshly prepared with acetate buffer (300 mM, pH=3.6), 10 mM 2,4,6-tripyridyl-s-triazine (TPTZ) solution, iron (III) chloride anhydrous solution (20 mM) at a ratio of 10:1:1 (v/v/v). The The above reconstituted methanol extracts were mixed with FRAP reagent and incubated in the dark at 37 °C for 5 min. The absorbance at 593 nm was measured using the Varioskan LUX Multimode Microplate Reader. The FRAP values were expressed as μmol Fe^2+^ equivalents per g dry weight of the freeze-dried extract (μmol Fe^2+^/g DW), based on the Fe^2+^ standard curve generated from serial dilutions of ferrous sulfate against a blank control.

#### 2.3.5. Characterisation of bioactive compounds in the ethanol extract via HPLC-ESI-QTOF-MS/MS

The ethanol extract of Kakadu plum was reconstituted and filtrated through 0.45 µm nylon syringe filters (Sigma-Aldrich, Australia) before dispensed into labelled LC vials. An Agilent 6520 Q-TOF LC/MS system coupled with Agilent 1200 Infinity Series LC (Agilent Technologies, Santa Clara, CA, USA) was employed to characterise the bioactive compounds according to Adiamo et al. (2024), with minor modification. The separation was achieved using a Synergi 4 µm Hydro-RP 80 Å LC column 100 x 2 mm protected by an AQ C18 guard column (4.0LJ×LJ3.0 mm) (Phenomenex, Lane Cove, NSW, Australia). The mobile phase A (0.1% formic acid in Milli-Q water) and mobile phase B (0.1% formic acid in acetonitrile) were delivered at 0.2 mL/min with the following gradient: 0-20 min (5-15 % B), 20-45 min (15-40% B), 45-60 min (40-60% B), 60-70 min (60-95% B), 70-80 min (95 % B), 80-80.1 min (95-5% B), and 80.1-85 min (5 % B). The LC condition was set as follows: column temperature 30 °C, injection volume 10 μL and flow rate 0.2 mL/min.

#### 2.3.6. Fourier transform infrared spectroscopy (FTIR) spectra analysis

The functional groups of the freeze-dried Kakadu plum water and ethanol extracts were determined by the Spectrum Two FT-IR spectrometer (PerkinElmer, Inc., Victoria, Australia). Spectra were collected over the wavenumber range of 400-4000 cm^-1^ with 32 scans at the spectral resolution of 4.0 cm^-1^ (Jia et al., 2021).

### 2.4. Network pharmacology analysis

A small molecule pharmacology workflow was used to link the identified compounds to oxidative-stress biology and to prioritise compound-target pairs for structural validation. The analysis was proceeded in five stages: 1. compound-target prediction and disease-oriented filtering; 2. Network pharmacology analysis including compound-target interaction, protein-protein interaction (PPI) network construction and functional enrichment analysis; 3. multi-stage ADMET and developability screening; 4. cross-species ortholog mapping and comparative validation; and 5. pharmacokinetic compartment screening followed by molecular docking of the most feasible pairs. A pipeline of the forementioned *in silico* analysis has been established by our group and will be made available on PangenomeAI platform https://github.com/PangenomeAI.

#### 2.4.1. Network pharmacology and target identification

Potential targets of the identified compounds were predicted with the SwissTargetPrediction web server (https://www.swisstargetprediction.ch/ restricted to Homo sapiens, probability ≥ 0.1) and processed using NetPharmPy (v1.0.0). In parallel, an oxidative-stress reference gene set was assembled from four databases (Gene Ontology [GO], WikiPathways, MSigDB and Reactome). The predicted compound targets were intersected with this reference set to define the shared oxidative-stress targets, from which a compound-target network was constructed. A protein-protein interaction network of the shared targets was built by mapping to the STRING database (v12.0; Homo sapiens; confidence score ≥ 0.7), and core targets were identified through two successive rounds of median-threshold screening on five topological metrics: degree centrality (DC), eigenvector centrality (EC), local average connectivity (LAC), betweenness centrality (BC) and network centrality (NC). Functional enrichment of the shared targets against GO terms (biological process, cellular component and molecular function) and KEGG and Reactome pathways was performed with g:Profiler (e110_eg57_p18_4b54045), applying a Benjamini-Hochberg false discovery rate (FDR) threshold of p < 0.05.

#### 2.4.2. Multi-stage ADMET and developability screening

The pharmacokinetic and toxicity profiles of the network-derived candidates were evaluated using an integrated ADMET screening funnel. Physicochemical properties and drug-likeness (Lipinski’s Rule of Five and Veber’s criteria) were calculated using RDKit (v2023.09.1). Absorption, Distribution, Metabolism, Excretion, and Toxicity (ADMET) properties were predicted using ADMET-AI (v1.0.0), a graph neural network-based predictor. Structural alerts for toxicity and pan-assay interference compounds (PAINS) were identified using the RDKit FilterCatalog. Mechanistic toxicokinetic parameters, including steady-state plasma concentration and total clearance, were estimated via the httk (v2.2.1) R package using a three-compartment model.

#### 2.4.3. Pharmacokinetic compartment screening and pair feasibility

To ensure physiological relevance, compound-target pairs were subjected to a route-specific compartment screening protocol using the pair_compartment_screening module developed by our group. The principles of screening are described as below: for the oral administration route, compounds were filtered based on a multi-parameter pharmacokinetic gate: Human Intestinal Absorption (HIA) > 30%, Caco-2 permeability (log Papp) > -5.15, and a preference for non-substrate status regarding P-glycoprotein (P-gp) efflux to ensure sufficient systemic exposure.

A compartment-matching algorithm was then applied to verify that the predicted systemic distribution of the compound aligned with the anatomical localization of the target protein. Target localization was retrieved from UniProt and the Human Protein Atlas, while compound accessibility was determined by predicted Blood-Brain Barrier (BBB) penetration (for CNS targets) and volume of distribution (V_ss_). Only “pharmacokinetically feasible” pairs, in which the compound demonstrated both high oral bioavailability and the ability to reach the target’s physiological compartment, were carried forward for structural validation.

#### 2.4.4. Cross-species ortholog mapping and comparative validation

To evaluate the evolutionary conservation of the identified compound-target interactions and support potential translational research, human core targets were mapped to their respective orthologs in the dog model organisms (i.g., *Canis lupus familiaris*) using the ortholog_mapping tool developed by our group. In principle, ortholog identification was executed via UniProtKB using organism-scoped gene-symbol matching, with a prioritised selection of reviewed (Swiss-Prot) entries to ensure high-quality reference sequences.

For each identified animal ortholog, three-dimensional protein structures were retrieved from the AlphaFold Protein Structure Database. In instances where high-confidence AlphaFold models were unavailable, experimental structures were sourced from the RCSB Protein Data Bank (PDB) as a fallback. These animal-specific structures were then used to assemble a comparative docking-ready pair list. This enabled a side-by-side assessment of binding affinities and interaction fingerprints between human and animal models, providing a computational basis for cross-species efficacy and safety evaluation.

#### 2.4.5. Molecular docking and interaction profiling

Molecular docking simulations were performed to quantify the binding affinity between the screened compounds and their respective targets. Protein structures were retrieved from the RCSB Protein Data Bank (PDB) or modelled via AlphaFold2. Ligands were prepared using Open Babel (v3.1.1) to assign proper protonation states at pH 7.4. Docking was executed using AutoDock Vina (v1.2.5) with an exhaustiveness setting of 8. The search space was defined by a grid box centred on the known active site or the geometric centre of the protein. Protein-ligand interaction fingerprints (PLIFs) were generated using ProLIF (v2.0.0) to identify key hydrogen bonds, hydrophobic contacts, and π-stacking interactions.

### 2.5. Intracellular antioxidant effect of the Kakadu plum extract against the H_2_O_2_-induced oxidative stress

#### 2.5.1. Cell culture

Canine primary small intestinal epithelial cells (# D-6051, Cell Biologics Inc., Chicago, USA) were immortalised using SV-40 virus, which was kindly provided by the Walter and Eliza Hall Institute (Parkville, VIC, Australia). The cells were cultured in filtered-cap T25 flasks with the complete epithelial cell medium (#M6621, Cell Biologics Inc., Chicago, USA) supplemented with 1% penicillin-streptomycin in a 37 C humidified incubator (BB 150 CO2 incubator, Thermo Fisher Scientific Australia Pty Ltd, VIC, Australia) with 5% CO_2_. The cells were observed every day under the FLoid™ Cell Imaging Station (Life Technologies Australia Pty Ltd., VIC, Australia) inverted microscope, and subculture was conducted when cells reached 80% confluence. A detailed evaluation of cell performance and morphology has been reported in a companion paper (He et al., in press).

#### 2.5.2. Cell cytotoxicity assay

To determine the non-cytotoxic concentrations of water and ethanol extracts, the Lonza ViaLight® Plus cell proliferation cytotoxicity assay was performed on the immortalised canine small intestinal cell line at different concentrations of the samples. The cells were seeded at 2×10^4^ per well in 96-well plates with antibiotic-free medium, and then were treated with Kakadu plum water extract (0-500 µg/mL) and ethanol extract (0-350 µg/mL) for 24 h. The ethanol extract was freeze dried and reconstituted to water prior to cell culture experiments to avoid cytotoxicity of ethanol. After 24 h, cell lysis reagent was added to each well for 10 min before adding the luminescent dye. The luminescence at wavelength 562 nm was measured using a NIVO plate reader (PerkinElmer, Inc., USA). The results were normalized to the untreated control as 100%.

#### 2.5.3. Generation kinetics of intracellular ROS

The *in vitro* oxidative stress cell model was established using H_2_O_2_, a well-known oxidative stress inducer (Gao et al., 2026; C. Li et al., 2023). To determine the dosages for the H_2_O_2_ on the immortalised canine small intestinal cell line, the cytotoxicity assay against cells treated with H_2_O_2_ (50-1800 µM) for 24 h was conducted according to Mohammed et al. (2023). The concentration of H_2_O_2_ at 80% cell viability and IC50 were used to induce mild and severe cellular oxidative stress for the subsequent experiment.

To determine the intracellular ROS generation in live cells over time, a cell-permeable non-fluorescent probe, CM-H2DCFDA (#C6827, ThermoFisher Scientific Australia Pty Ltd, VIC, Australia) was employed. Cells were seeded in dark, clear-bottom 96-well microplates at a density of 2 ×10^4^ cells/well incubating under the standard conditions for 24 h. Culture medium was then replaced with 5 μM of CM-H2DCFDA solution dissolved in HBSS without phenol red and incubated in the dark under standard conditions for 30 min. After incubation, the CM-H2DCFDA solution was removed, and cells were gently washed twice with HBSS before the addition of Kakadu plum extracts. Intracellular ROS generation over time was determined by measuring CM-DCF fluorescence intensity at 0, 15, 30, 45 min, and 1 h, followed by every 30 min for up to 8 h. Fluorescence was monitored at excitation and emission wavelengths of 495/20 nm and 530/30 nm, respectively, using the NIVO plate reader. ROS production was expressed as the fold change in fluorescence intensity relative to the untreated control, where the control at 0 h was set to 1.

### 2.6. Statistical analysis

Statistical analysis was performed using the SPSS Statistics for Windows, version 27.0 (IBM Corp., Armonk, NY, USA), the GraphPad Prism for Windows, version 9.1 (Dotmatics Ltd, Boston, MA, USA), and the Agilent LC-ESI-QTOF-MS/MS MassHunter Workstation software, version B.03.01 (Agilent Technologies, Inc., Santa Clara, CA, USA). All groups were analysed in technical triplicate (n=3). The one-way analysis of variance (ANOVA) followed by Tukey’s post hoc test at a 95% confidence interval was employed to assess significant (p < 0.05) differences.

## 3. Results and Discussion

### 3.1 Physicochemical properties of Kakadu plum extract

#### 3.1.1 Chemical antioxidant assay

To determine the effect of extraction solvents and extraction technologies on the physicochemical characteristics of Kakadu plum, the freeze-dried KP fruit powder was extracted using different solvents assisted with different processing technologies.

Figure 1 revealed that the extraction solvent was the dominant factor influencing chemical antioxidant properties, with no effects of extraction technique or solvent–technique interaction (p > 0.05). This limited impact of extraction technique may be attributed to the exceptionally fine particle size of the Kakadu plum powder (filtered through a 1000-mesh screen), as milling-induced cell wall disruption facilitates efficient mass transfer and near-complete release of bioactive compounds. Smaller particle sizes have been shown to enhance the antioxidant capacity of plant extracts like tea and ginger (Makanjuola, 2017), and the thorough release of phenolics likely renders additional extraction techniques redundant. Therefore, subsequent analysis focused on comparing antioxidant capacity across the three extraction solvents.

**Figure 1.**
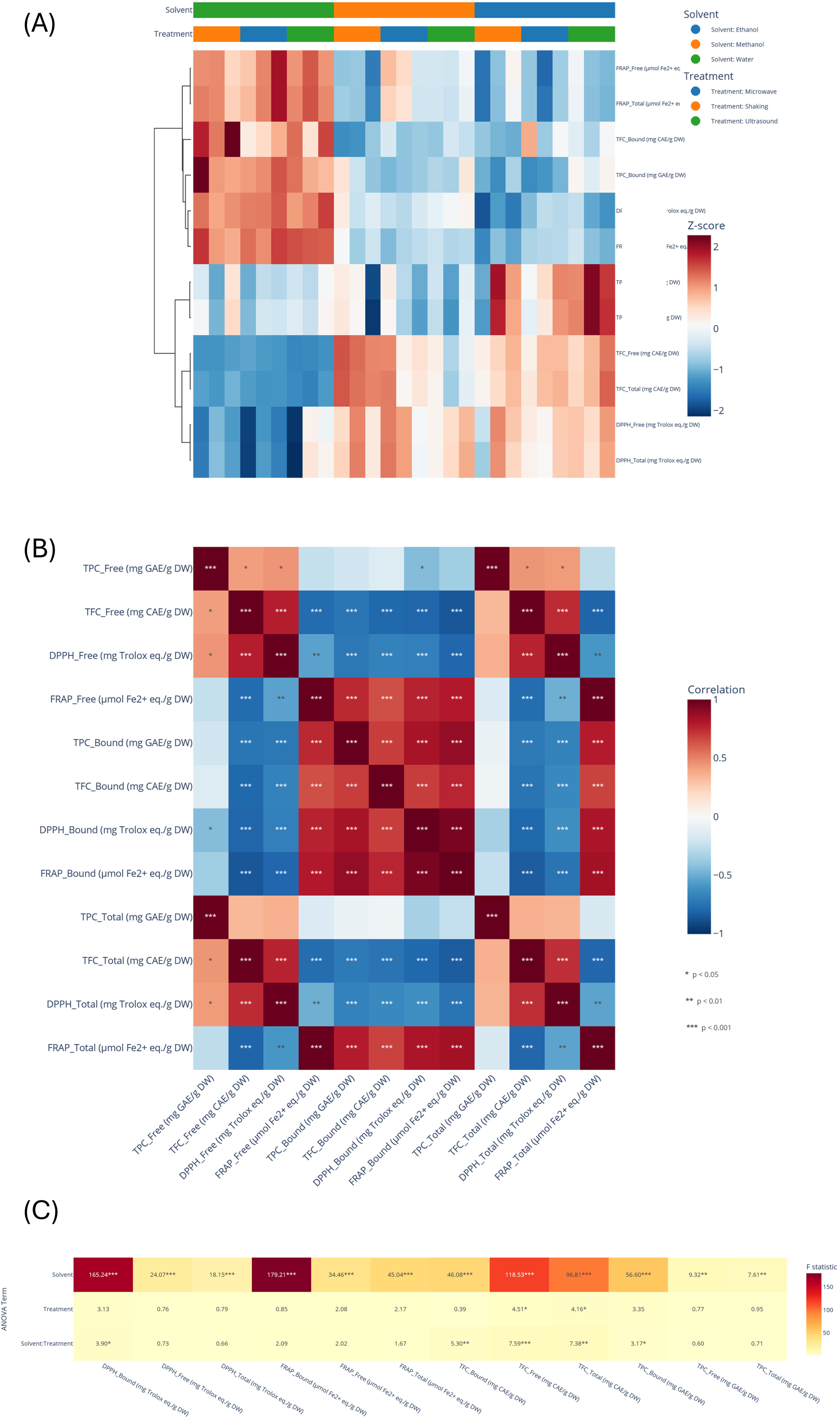
Effect of extraction solvents and technologies on the antioxidant properties (i.e., TPC, TFC, DPPH, and FRAP) for free, bound, and total phenolics of Kakadu extracts. A: Heatmap of the antioxidant properties (i.e., TPC, TFC, DPPH, and FRAP) of Kakadu plum extracts across different extraction solvent and technologies. B: Correlation matrix of antioxidant properties (i.e., TPC, TFC, DPPH, and FRAP) for free, bound, and total phenolics of Kakadu plum. C: Multi-way ANOVA heatmap of the effects of extraction solvents and technologies on antioxidant properties (i.e., TPC, TFC, DPPH, and FRAP). Data represents the mean of triplicates. Statistical significance is given as follows: * p < 0.05, ** p < 0.001, and *** p < 0.005, as compared to untreated controls. TPC, total phenolic content, mg GAE/ g DW; TFC, total flavonoid content, mg CAE /g DW; DPPH, 2,2-diphenyl-1-picrylhydrazyl, mg TE/ g DW; FRAP, ferric reducing antioxidant power, μmol Fe2+/g DW.

The TPC, TFC, DPPH, and FRAP for free, bound, and total phenolics of Kakadu extracts using different solvents (i.e., ethanol, methanol, and water) were visualised in Figure 2 and values were listed in Table S1. For TPC, the free phenolics extracted using ethanol performed the best, at 225.3 mg GAE/g DW, significantly higher than the methanol extract 204.8 mg GAE/g DW and water extract 205.7 mg GAE/g DW (Table S1). The TPC values of free phenolics of Kakadu plum were in line with the values range of 200.2 to 382.5 mg GAE/g DW by Konczak et al. (2014) and 198.4 to 240.3 mg GAE/g DW obtained using 70% ethanol by J. Li et al. (2023). The TPC of total phenolics showed a similar trend to the free phenolic fraction. The TFC values of free and total phenolics fraction in Kakadu plum extracts using ethanol and methanol were significantly higher than those of the water extract. For free and total phenolics (Figure 2B and 2J), the TFC values of ethanol (6.1 mg CAE/g DW) and methanol (5.9 mg CAE/g DW) extracts were more than 2-fold greater than those of water extracts (2.6 mg CAE/g DW). No significant difference (p > 0.05) was observed between the ethanol and methanol groups with respect to TFC values, indicating that ethanol and methanol exhibited comparable extraction efficiency for flavonoid compounds. In previous studies, the TFC of Kakadu plum was measured using the quercetin equivalents (QE) at the range of 1.1 to 1.3 mg QE /g DW (Akter et al., 2021) and 1.4 mg QE /g DW extracted using 70% ethanol (J. Li et al., 2023). The DPPH showed a very similar trend to that of the TFC (Figure 2C, 2G, and 2K). The DPPH of free and total phenolic fractions in the water extracts was significantly (p<0.001) lower than those of the methanol and ethanol groups. For the free and total phenolic fraction, samples extracted using ethanol (259.22 mg Trolox equivalents (TE)/g DW) and methanol (264.1 mg TE/g DW) showed similar performance (p > 0.05), which was comparable to the DPPH values reported in a previous study using methanol-assisted ultrasound extraction (215.9 to 239.7 mg TE/g DW) (Akter et al., 2021). Figures 2D, 2H, and 2L illustrated the FRAP values of the Kakadu plum extracts, indicating comparable extraction efficiency between methanol and ethanol groups for FRAP values (p > 0.05). Water extract exhibited the highest FRAP value (2202.0 μmol Fe^2+^ eq./g DW), which was slightly lower than the range reported in the previous research (2800.9 to 4197.5 μmol Fe^2+^ eq./g DW) (Konczak et al., 2014). This could be attributed to differences in sample origin, maturity condition, or extraction method.

**Figure 2.**
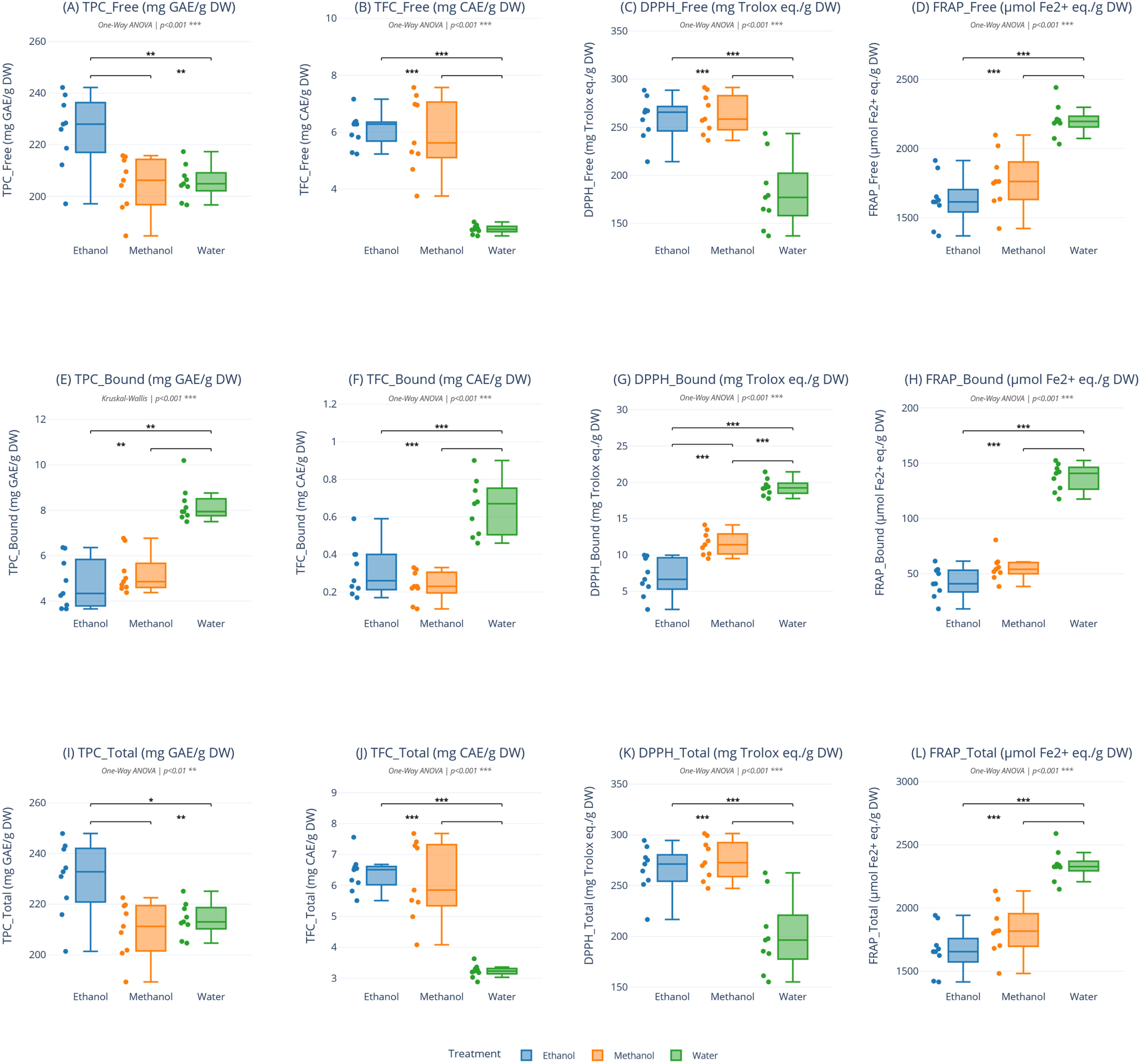
The antioxidant properties (i.e., TPC, TFC, DPPH, and FRAP) for free, bound, and total phenolics of Kakadu extracts using different solvents. A: TPC of free phenolics of Kakadu extracts. B: TFC of free phenolics of Kakadu extracts. C: DPPH of free phenolics of Kakadu extracts. D: FRAP of free phenolics of Kakadu extracts. E: TPC of bound phenolics of Kakadu extracts. F: TFC of bound phenolics of Kakadu extracts. G: DPPH of bound phenolics of Kakadu extracts. H: FRAP of bound phenolics of Kakadu extracts. I: TPC of total phenolics of Kakadu extracts. J: TFC of total phenolics of Kakadu extracts. K: DPPH of total phenolics of Kakadu extracts. L: FRAP of total phenolics of Kakadu extracts. Data represents the mean of triplicates. Statistical significance is given as follows: * p < 0.05, ** p < 0.001, and *** p < 0.005, as compared to untreated controls. TPC, total phenolic content, mg GAE/ g DW; TFC, total flavonoid content, mg CAE /g DW; DPPH, 2,2-diphenyl-1-picrylhydrazyl, mg TE/ g DW; FRAP, ferric reducing antioxidant power, μmol Fe2+/g DW

The higher FRAP of the water extract’s free phenolic fraction, despite its lower TPC, TFC and DPPH, may arise from the differing chemistry of each assay combined with solvent selectivity. TPC and TFC quantify phenolic content rather than reducing capacity. FRAP measures ferric reduction via single-electron transfer, whereas DPPH scavenging proceeds mainly by hydrogen-atom transfer, which may diverge when different compound classes dominate (Chen et al., 2020; Hossain et al., 2021). Those small hydrophilic reductants as potent electron donors are abundant in Kakadu plum (e.g., ascorbic, ellagic and gallic acids) and are more preferentially extracted by water (He et al., 2023). The less polar ethanol and methanol recover more flavonoids and proanthocyanidins, lifting the TPC and TFC; the B-ring hydroxyls in the organic solvents are effective hydrogen donors that favour DPPH antioxidant activity, however, are weak electron donors, does not increase FRAP proportionally (Lang et al., 2024). The water fraction’s higher share of bound phenolics, reflecting the lower aqueous solubility of free phenolics, further explains why FRAP decouples from TPC, TFC and DPPH here (Makanjuola, 2017).

#### 3.1.2 Characterization of phytochemical profile of Kakadu plum via HPLC-Q-TOF-MS/MS

The composition of phenolic compounds for the ethanol extract of Kakadu plum was revealed using HPLC-ESI-QTOF-MS/MS, a total of 26 compounds were identified (Table 1), which were dominated by hydrolysable tannins and triterpenoids. Of the 26 compounds, 24 phenolic constituents (excluding ascorbic acid and α-linolenic acid) were carried forward to the network pharmacology analysis. Four organic acids, including ascorbic acid, gallic acid, chebulic acid, and monogalloylshikimic acid were identified. Ascorbic acid (m/z 175.0234 [M−H]C) was detected at a retention time of 1.639 min, producing characteristic fragment ions at m/z 115 and 87, corresponding to the sequential loss of COC and HCO moieties. The early elution of ascorbic acid and gallic acid is attributable to their high polarity and align with previous studies of Konczak et al. (2014) and Akter et al. (2019). Chebulic acid (m/z 355.0650 [M−H]C) displayed characteristic fragments at m/z 265, 295, and 337, consistent with reported fragmentation behaviour in *Terminalia* species (Pfundstein et al., 2010). Hydrolysable tannins constituted the principal polyphenolic compound class, with 11 compounds identified, which agrees with previous studies (Phan et al., 2022). Ellagic acid (m/z 300.9970 [M−H]C), one of the most abundant phenolic compounds in Kakadu plum (Konczak et al., 2014), was identified. Ellagitannins, such as chebulagic acid (m/z 953.0836 [M−H]C) and chebulinic acid (m/z 955.0993 [M−H]C), were identified according to the diagnostic fragment ions at m/z 301 (ellagic acid moiety) and m/z 337 (galloylshikimic acid derivative), supported with previous studies in *T. ferdinandiana* (Bobasa et al., 2022) and *T. chebula (*Pfundstein et al., 2010). The identification of 1,3,6-tri-O-galloyl-β-D-glucose (m/z 635.0849 [M−H]C) and tetragalloylglucose (m/z 787.0939 [M−H]C) indicates the presence of a gallotannin biosynthetic series in Kakadu plum. The fragmentation of tetragalloylglucose produced sequential losses of galloyl units (m/z 787 → 635 → 465), consistent with the progressive de-galloylation pattern reported for gallotannins (Akter et al., 2019). Tellimagrandin I (m/z 785.0795 [M−H]C) was also detected with fragments at m/z 301 (ellagic acid) and m/z 617 (loss of gallic acid). Notably, several methylated ellagic acid derivatives were identified, including 3,3′-di-O-methylellagic acid 4-O-xyloside, trimethylellagic acid glucoside, and 3,3′,4′-tri-O-methylellagic acid (Landete, 2011). Two flavone C-glycosides were identified, luteolin-8-C-glucoside (447.0905 [M−H]C) and isovitexin (apigenin-6-C-glucoside, m/z 431.0959 [M−H]C). Eight triterpenoids were also identified. Arjungenin (m/z 503.3354 [M−H]C), arjunolic acid (m/z 487.3405 [M−H]C), along with their corresponding glycosides, arjunglucoside I (m/z 711.3916 [M−H]C) and arjunolic acid hexoside (m/z 695.3974 [M−H]C), are well-characterised constituents of the genus *Terminalia*, particularly *T. arjuna*, where arjunolic acid has been reported as a major bioactive compounds (Ghosh & Sil, 2013). Asiatic acid (m/z 487.3401 [M−H]C) and madecassic acid (m/z 503.3346 [M−H]C) are ursane-type triterpenoids more commonly associated with *Centella asiatica* but have also been reported in *Terminalia* species (Cock, 2015). Linolenic acid (peak 24, m/z 277.2160 [M−H]C) was identified as the only fatty acid, which was reported in Kakadu plum kernel oil (Akter et al., 2018).

**Table 1.**
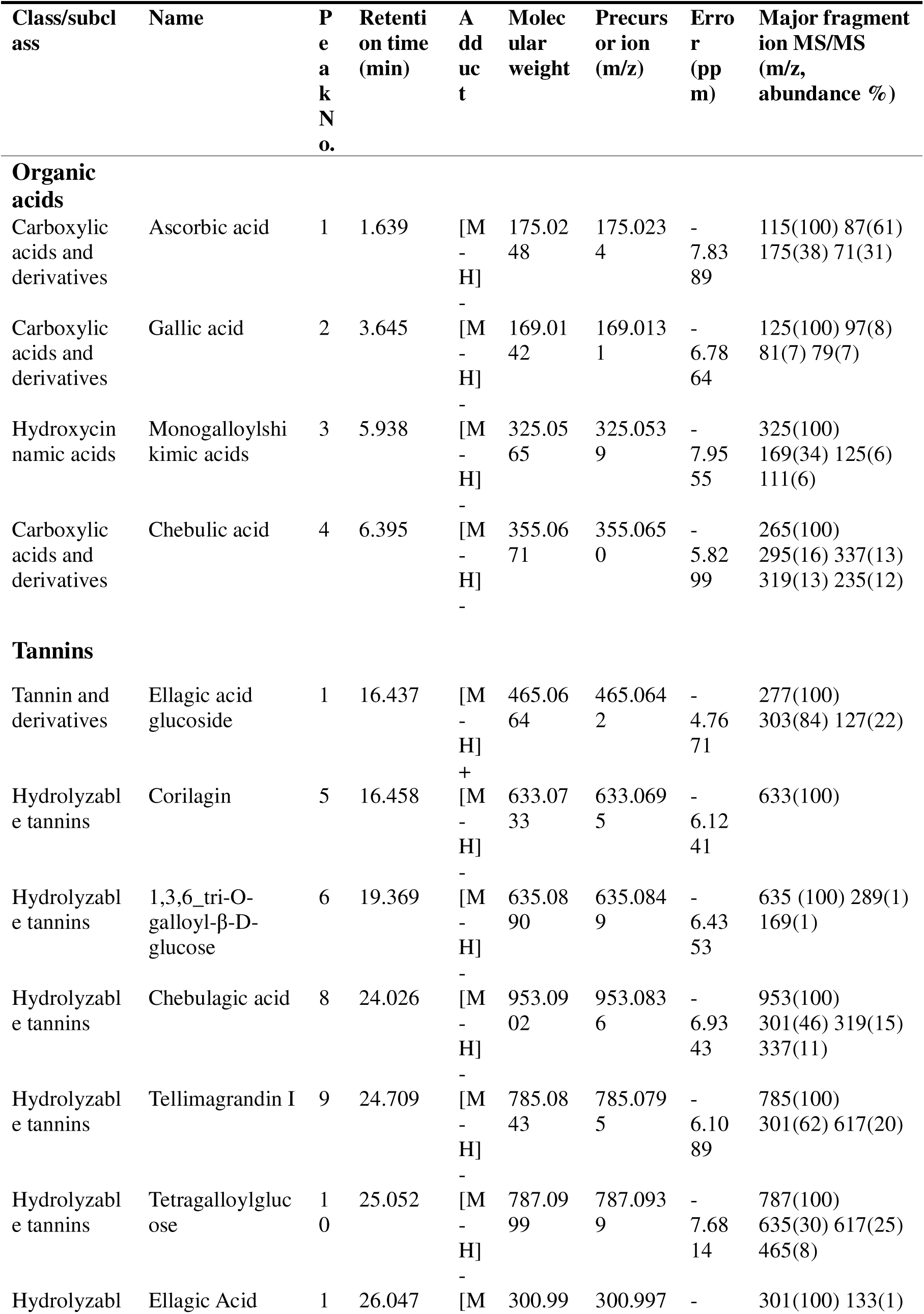
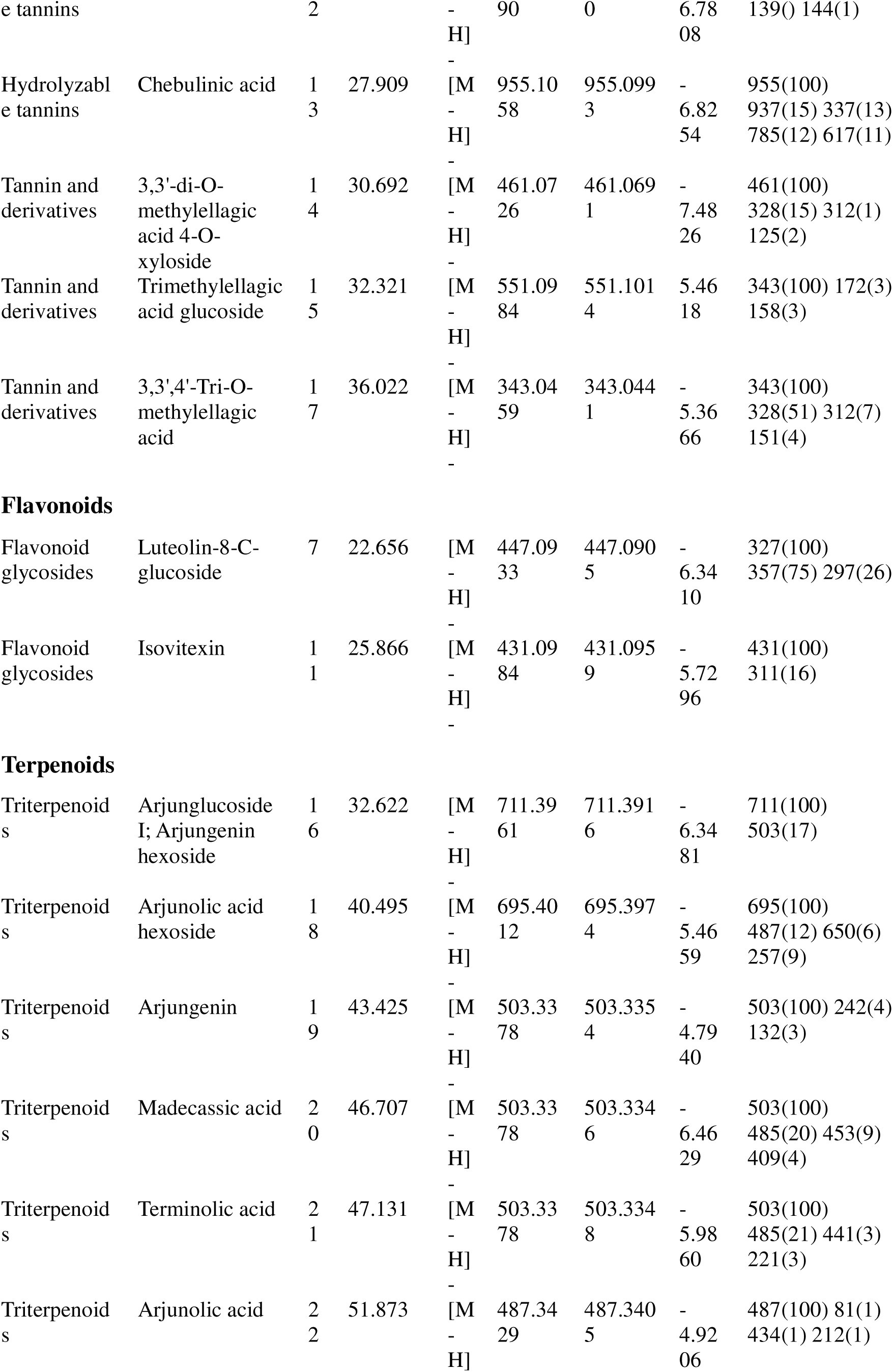
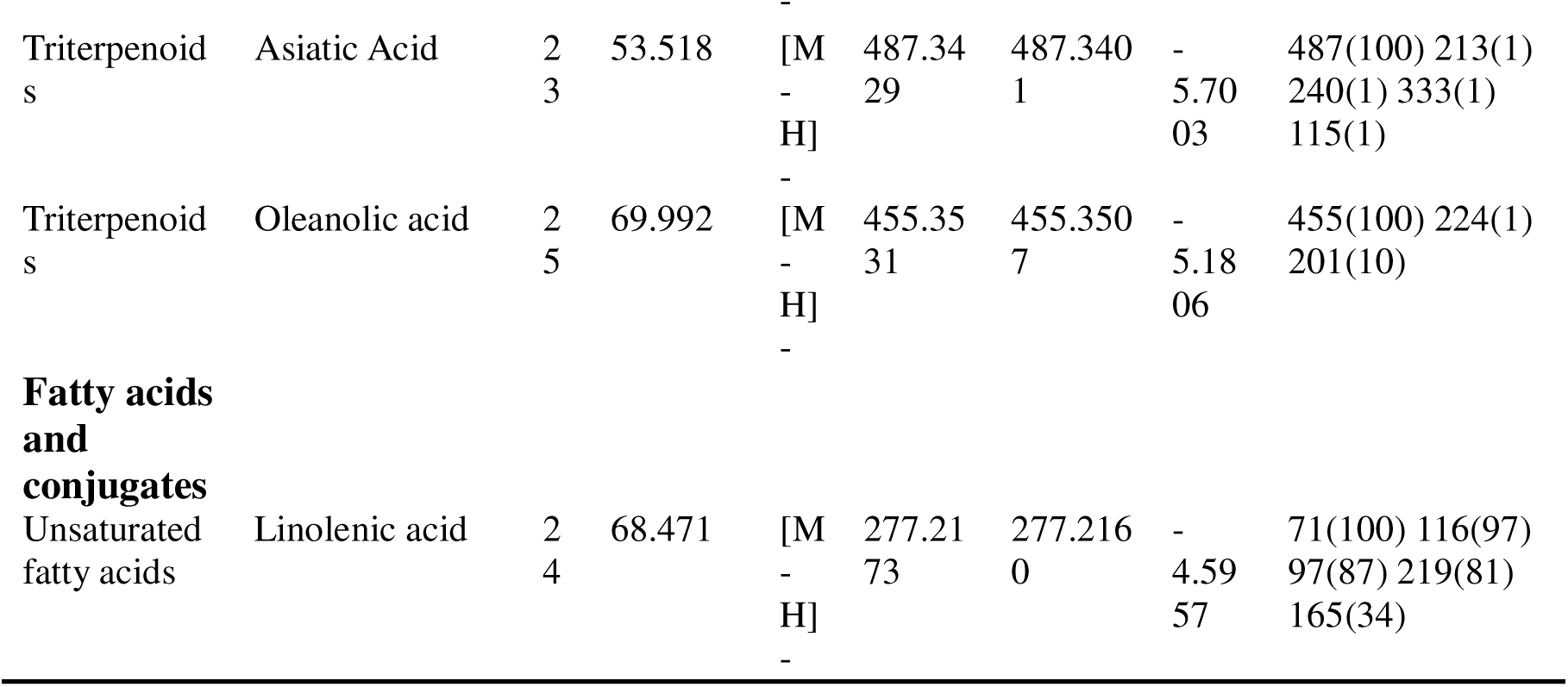
Identification of phenolic compounds for the ethanol extract of Kakadu plum using HPLC-ESI-QTOF-MS/MS.

FT-IR spectroscopy was performed to characterise the functional groups in both water and ethanol extract powders of Kakadu plum, with spectra acquired in the wavenumber range of 400-4000 cm-1 (Figure S3). There are two major peaks observed between the regions 3600 and 2800 cm^-1^. The peak at 3304 cm^-1^ resulted from the O-H stretching of Kakadu plum powder and inter- and intra-molecular hydrogen bonding of the galacturonic acid backbone of the molecule. The second peak observed around 2925 cm^-1^ corresponds to C-H absorption including CH, CH2 and CH3 stretching and bending vibrations (Khaligh & Asoodeh, 2022). According to Ramadhania et al. (2022), the transmitted peaks at 2910 and 1710 cm^-1^, corresponding to C-H and C=O (ketones) bonds, were associated with the presence of aromatic compounds. For the region between 1800-1500 cm^-1^, the peak at 1682 cm^-1^ corresponds to the C=O groups. Transmission peaks observed at 1201 were proposed to be contributed by the stretching of C-O bonds of carbonyl groups or C-O of alcoholic groups and glycosidic linkages of the phytochemical constituents of Kakadu plum. The region between 400 and 1200 cm^-1^ is considered the ‘fingerprint area’. The peak at 1031 cm^-1^ was detected in the spectrum, which aligns with the 1019 cm^-1^ being observed for Kakadu plum powder and pectin in previous analysis by Chaliha (2018).

### 3.2 Key antioxidant prediction and potential action mechanisms in human physiology

To explore the potential contributions of the phenolic components of Kakadu plum in the intracellular antioxidant properties and suppressive activities in the cellular oxidative stress, the 24 phenolic compounds characterised in the extract were analysed by network pharmacology, integrated ADMET profiling, cross-species ortholog mapping, and molecular docking.

#### 3.2.1 Network pharmacology analysis

Potential oxidative-stress targets of the Kakadu plum phenolic compounds were first predicted with SwissTargetPrediction, and an oxidative-stress reference gene set was assembled from four curated databases (GO, WikiPathways, MSigDB and Reactome). The four sources contributed a non-redundant reference set of 766 genes (Figure 3A), with GO providing the largest share, confirming a broad, database-independent definition of oxidative-stress biology. Intersection analysis between predicted targets of Kakadu plum and ROS-related targets retrieved from the databases revealed 30 overlapping targets between the 196 unique human targets for the compound set and the 766 targets in the reference set (Figure 3B). These 30 shared targets define the mechanistic search space linking Kakadu plum chemistry to redox regulation, and a similar multiple-compound, multiple-target profile has been reported for other polyphenol-rich extracts (Hopkins, 2008; Merecz-Sadowska et al., 2025). Figure 3C visualised the compound-target network of 17 compounds and 30 selected targets, in which each compound engaged at least one oxidative-stress target, indicating that antioxidant activity is distributed across the extract rather than confined to one or two constituents.

**Figure 3.**
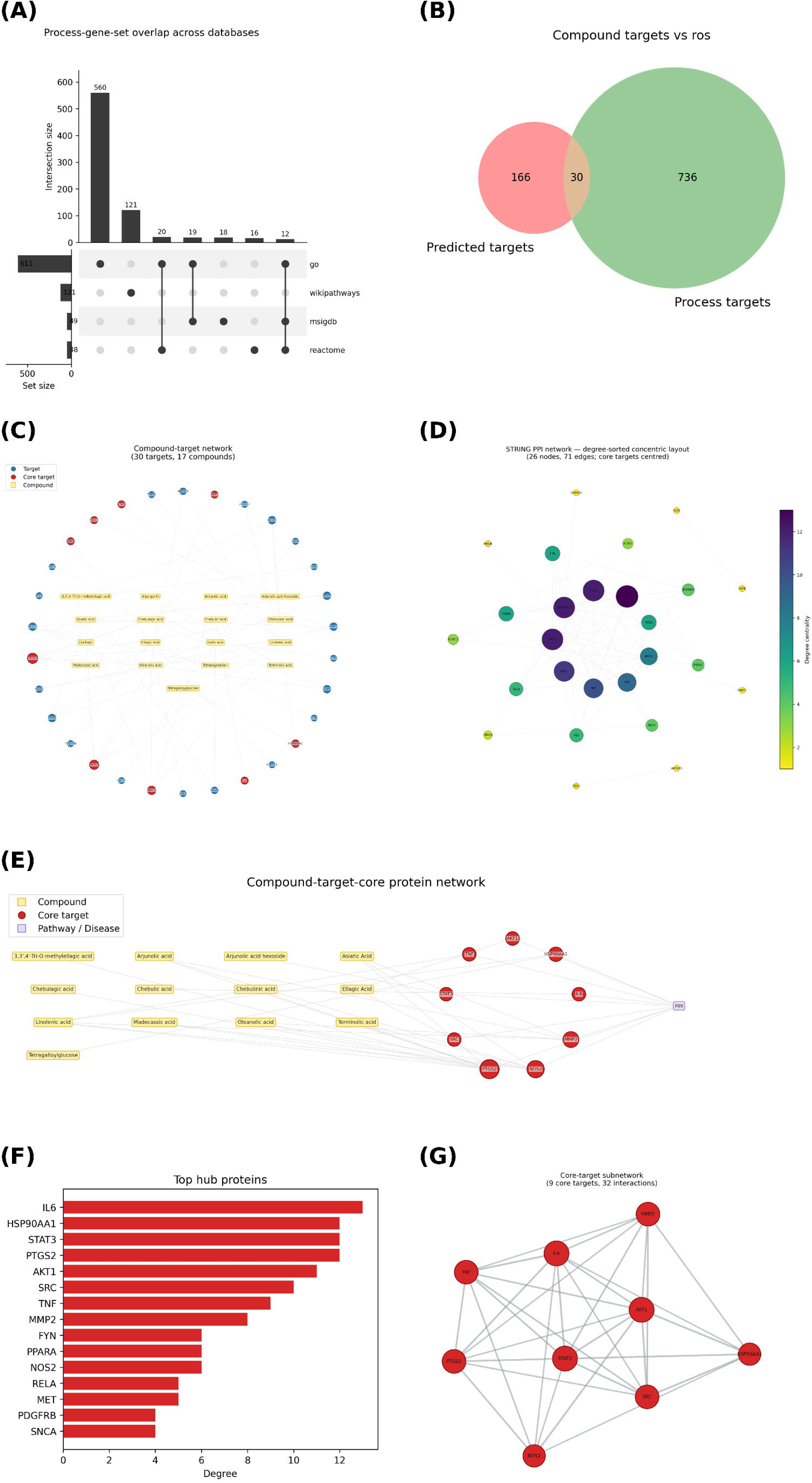
Target identification and network pharmacology overview. (A) Distribution of biological-process gene-set members across the queried databases (UpSet plot). (B) Overlap between predicted compound targets and biological-process genes (Venn diagram). (C) Compound–target network (targets on a circle, compounds in rows). (D) STRING PPI network (degree-sorted concentric layout; core targets centred, nodes coloured/sized by degree centrality). (E) Compound–core target–pathway/disease interaction network. (F) Top hub targets ranked by degree centrality. (G) Core-target protein–protein interaction subnetwork. Panels are shown in analysis order; database-overlap and Venn panels depend on the selected analysis mode.

Protein-protein interaction (PPI) analysis of the shared targets in STRING further constructed a network of 26 nodes and 71 edges (Figure 3D), indicating co-regulation across multiple oxidative-stress-related pathways. Ranking nodes by degree centrality identified IL6, HSP90AA1, STAT3, PTGS2, AKT1, SRC, TNF and MMP2 as the most connected proteins, followed by FYN, PPARA and NOS2 (Figure 3F). Nine core target proteins were ultimately identified through two successive rounds of median threshold-based screening according to their degree centrality (DC), betweenness centrality (BC), eigenvector centrality (EC), network centrality (NC), and local average centrality (LAC) values as listed in Table 2 and their interactions were visualised in Figure 3G. Targets with DC, EC, LAC, BC, and NC values higher than 6, 0.2192, 4, 2.3708, and 6.4, respectively, were considered core targets, including IL6, PTGS2, HSP90AA1, STAT3, AKT1, SRC, TNF-alpha, MMP2 and NOS2. The compound-core target-disease network confirmed that multiple constituents converge on these nine proteins to reach the ROS phenotype (Figure 3E). Phenolic compounds may exert their biological effects in oxidative stress-related diseases through simultaneous regulation of multiple signalling pathways. PTGS2 (cyclo-oxygenase-2) and NOS2 (inducible nitric oxide synthase) are direct enzymatic sources of pro-oxidant and pro-inflammatory mediators, while IL6, TNF and STAT3 form a self-amplifying inflammatory circuit that both responds to and sustains ROS accumulation (Reuter et al., 2010). AKT1 and SRC are redox-sensitive kinases that transduce oxidative signals into survival and barrier-regulating responses, and HSP90AA1 is a stress-inducible chaperone central to the proteostatic response to oxidative injury (Corcoran & Cotter, 2013; Kumar et al., 2026). The recovery of this inflammation-redox module, rather than a set of isolated radical-scavenging enzymes, indicates that Kakadu plum phytochemicals attenuate epithelial oxidative stress at least partly by modulating an inflammatory signalling hub.

**Table 2.**
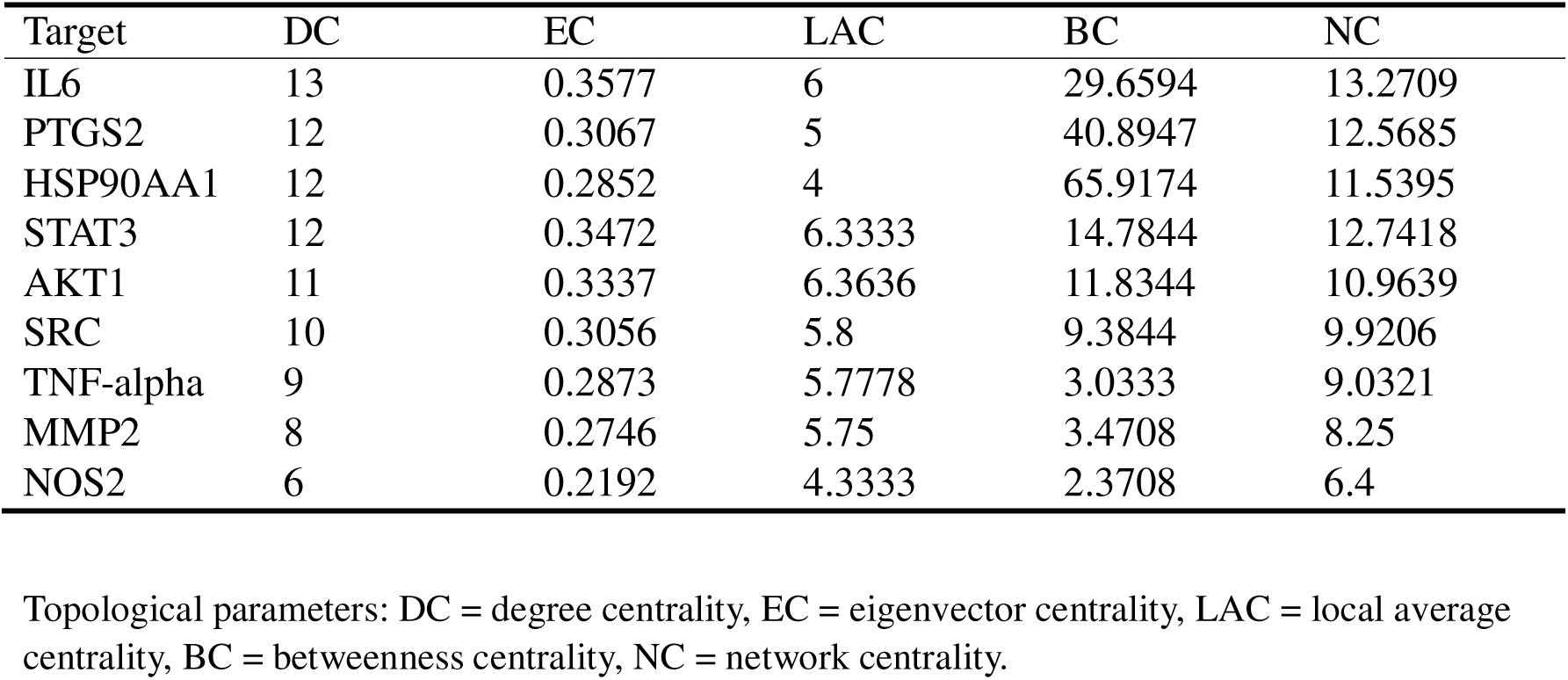
Network topological properties of core targets linked to oxidative stress-related diseases modulated by phenolic compounds in Kakadu plum.

| Target | DC | EC | LAC | BC | NC |
| --- | --- | --- | --- | --- | --- |
| IL6 | 13 | 0.3577 | 6 | 29.6594 | 13.2709 |
| PTGS2 | 12 | 0.3067 | 5 | 40.8947 | 12.5685 |
| HSP90AA1 | 12 | 0.2852 | 4 | 65.9174 | 11.5395 |
| STAT3 | 12 | 0.3472 | 6.3333 | 14.7844 | 12.7418 |
| AKT1 | 11 | 0.3337 | 6.3636 | 11.8344 | 10.9639 |
| SRC | 10 | 0.3056 | 5.8 | 9.3844 | 9.9206 |
| TNF-alpha | 9 | 0.2873 | 5.7778 | 3.0333 | 9.0321 |
| MMP2 | 8 | 0.2746 | 5.75 | 3.4708 | 8.25 |
| NOS2 | 6 | 0.2192 | 4.3333 | 2.3708 | 6.4 |

Topological parameters: DC = degree centrality, EC = eigenvector centrality, LAC = local average centrality, BC = betweenness centrality, NC = network centrality.

Functional enrichment of the 30 shared targets validated the target-identification step and the biology involved (Figure 4). Gene Ontology (GO) comprised three categories: biological-process, cellular-component, and molecular-function, with the top 14 terms in each category according to -Log10 Q value (Figure 4A). The biological-process terms were predominantly associated with response to oxidative and chemical stress. Cellular-component terms localised the targets to the cytoplasm, cytosol, cell periphery, plasma membrane, and vesicles, compartments central to ROS generation and handling. Molecular-function terms emphasised binding and catalytic activities, such as small molecule binding, ion binding, catalytic activity, identical protein binding, and oxidoreductase activity. KEGG enrichment was led by pathways in cancer, lipid and atherosclerosis, toxoplasmosis, and arachidonic acid metabolism (Figure 4B), associated with chronic oxidative and inflammatory stress. REAC enrichment converged on cytokine and interleukin signalling (including interleukin-4/interleukin-13), arachidonate metabolism, and synthesis of prostaglandins (Figure 4C). The recurrence of the arachidonic acid-prostaglandin axis across GO, KEGG and REAC maps onto the PTGS2 core target and onto the polyunsaturated fatty acid constituents of the extract, tying network topology, enrichment output and extract chemistry into one consistent account of intestinal redox and inflammatory control. These potential impacts of Kakadu constitute on biological targets were established on the condition that these compounds can reach the relevant biological locations of these targets.

**Figure 4.**
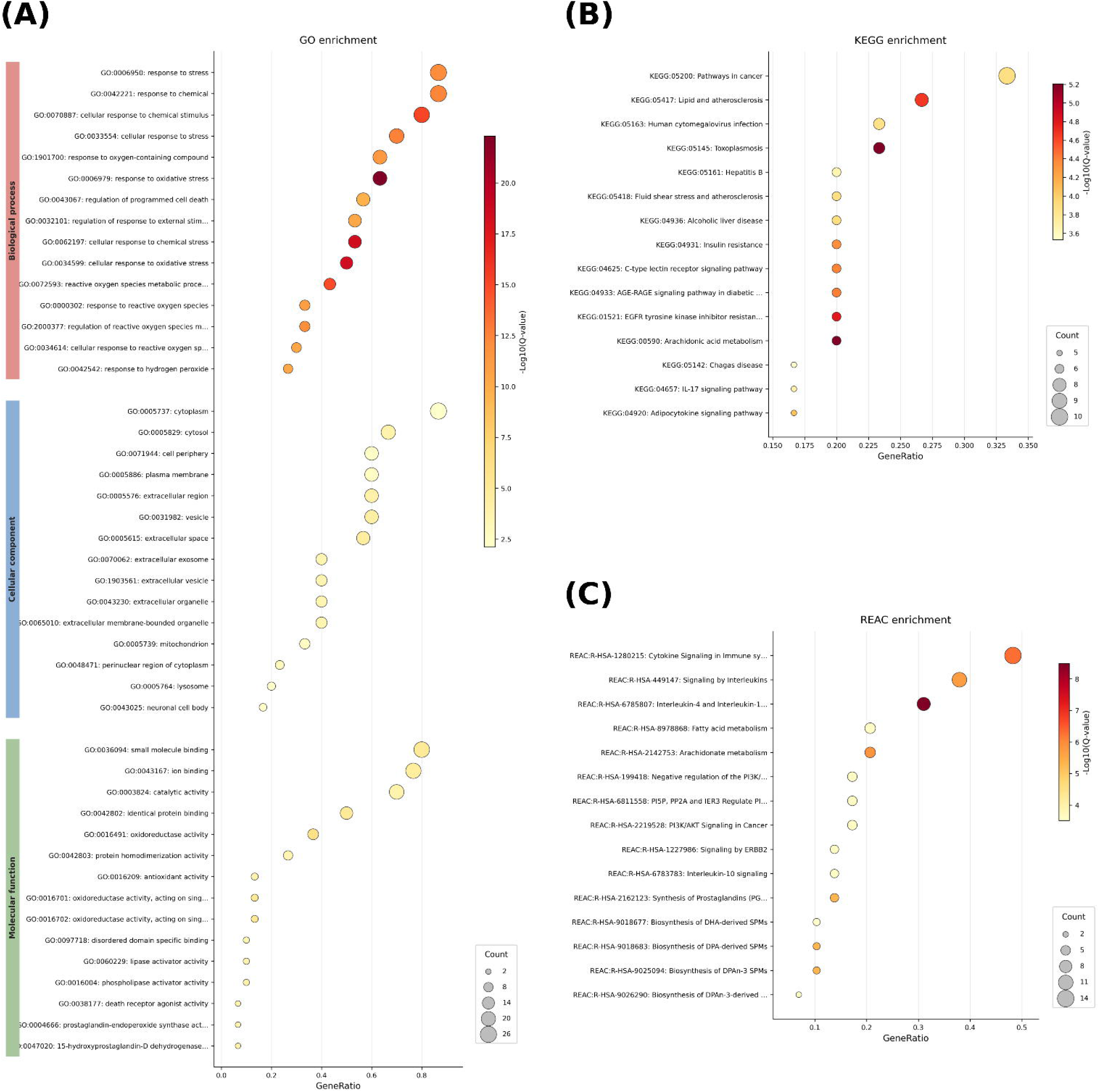
Functional enrichment analysis of the overlapping target set. (A) Gene Ontology enrichment (Biological Process, Cellular Component, Molecular Function). (B) KEGG pathway enrichment. (C) Reactome pathway enrichment. Dotplots show GeneRatio (x-axis), gene count (dot size) and −log10(adjusted P-value) (dot colour).

#### 3.2.2 ADMET screening

To screen the chemical compounds that could be used in pharmacology and drug discovery, the ADMET (absorption, distribution, metabolism, excretion, and toxicity) process was conducted. In total, 24 constituents passed through the integrated ADMET screen (Figure 5). The composite developability ranking was presented (Figure 5A), scored according to a weighted blend of QED, drug-likeness rule compliance, structural-alert burden and predicted oral bioavailability, with 3,3’,4’-tri-O-methylellagic acid, madecassic acid, terminolic acid, arjungenin, asiatic acid and arjunolic acid occupying the top positions, followed by ascorbic acid, gallic acid and ellagic acid. The large hydrolysable tannins (chebulinic acid, chebulagic acid, tellimagrandin I, tetragalloylglucose, corilagin) and glycosides (ellagic acid and arjunolic acid glycosides) ranked the lowest. Figure 5B evaluate compounds against drug-likeness rules. The top-ranked compounds satisfied the Lipinski, Veber and Egan rules consistent with the results in Figure 5A. The high-molecular-weight tannins violated multiple rules simultaneously. For an epithelial target acting on intestinal lumen and the apical membrane, poorly absorbed tannins still deliver antioxidant protection at the mucosal surface without requiring systemic uptake, while the readily absorbed triterpenoids and small phenolics could additionally contribute intracellular and post-absorptive effects. The structural-alert burden was assessed primarily according to the Dundee, MLSMR and PAINS filters (Figure 5C), suggesting that chebulinic acid, chebulagic acid, tellimagrandin I, tetragalloylglucose, corilagin and linolenic acid could be moderately hazardous. Figure 5D plotted the ADMET properties of 24 chemicals against the ADMET properties library developed by our group using USDA-Duke & MPD3 medical plant chemicals (n=8,272 known compounds) and ADMET-AI (v1.0.0). Chemicals located in the overlap between the favourable region of high predicted human intestinal absorption (above 0.9) and that of low predicted clinical toxicity (below 0.2) included ellagic acid, 3,3’,4’-tri-O-methylellagic acid, linolenic acid, madecassic acid, arjunolic acid, arjungenin, asiatic acid, and terminolic acid. Overall, the ADMET screen indicated that the triterpenoid and simple-phenolic fraction were the most developable systemic contributors and the hydrolysable tannins as luminally antioxidants, which could be referenced for the barrier-protective phenotype in the canine intestinal epithelial cell model.

**Figure 5.**
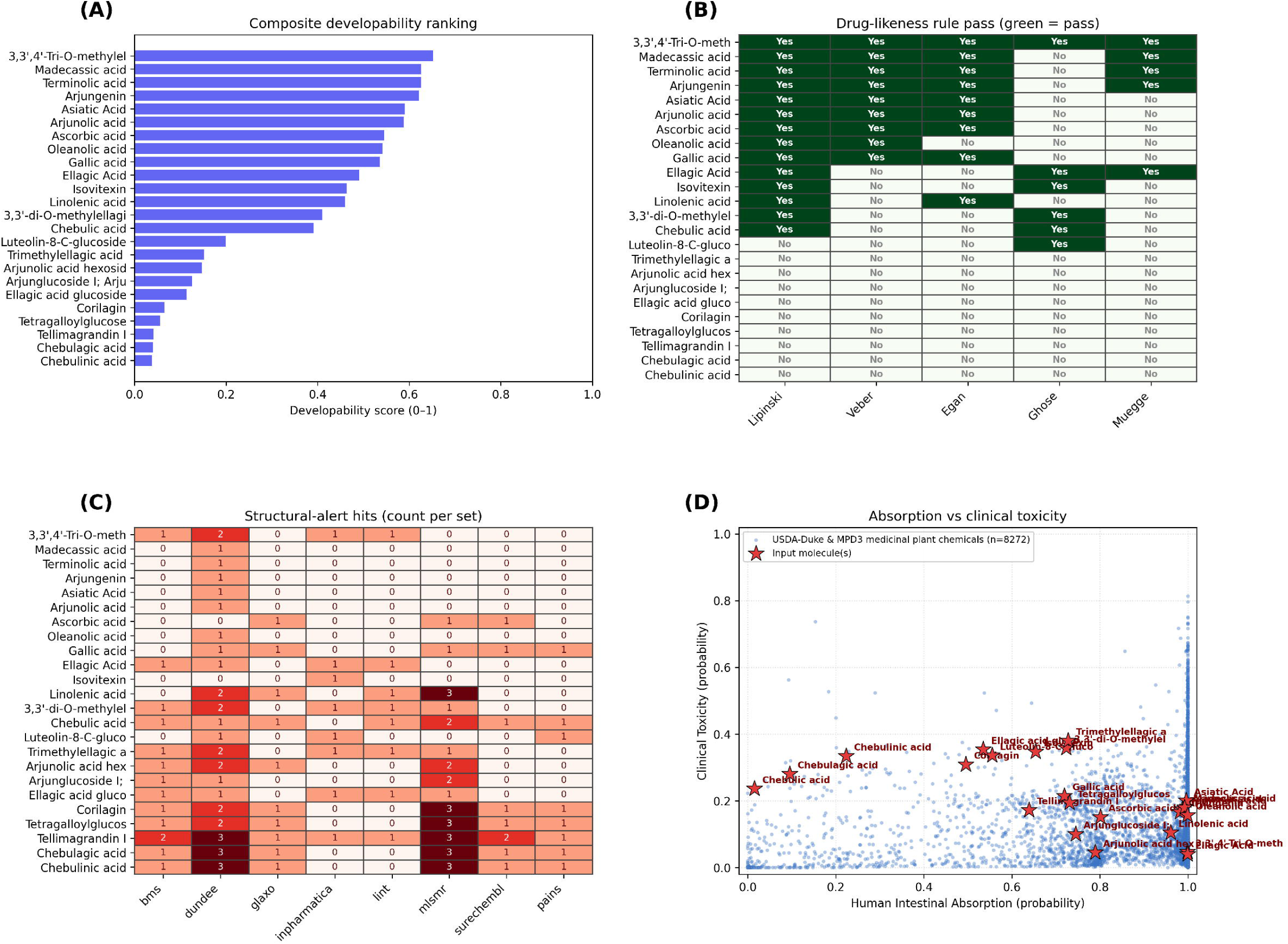
Integrated ADMET screening for 24 compounds identified in Kakadu plum ethanol extract. Composite developability ranking. Compounds ranked by the composite developability score (0–1; weighted blend of QED, drug-likeness rule compliance, structural-alert burden and predicted oral bioavailability). Longer bars indicate more favourable overall developability. (A) Drug-likeness rule compliance. Pass (green, “Yes”) / fail (pale, “No”) status for the Lipinski, Veber, Egan, Ghose and Muegge rule sets; each cell is annotated and gridded for clarity. (B) Structural-alert burden. Number of matched structural alerts per ChEMBL alert set (BMS, Dundee, Glaxo, Inpharmatica, LINT, MLSMR, SureChEMBL, PAINS); darker cells and higher counts indicate greater liability. (C) ADMET context plot. Predicted Human Intestinal Absorption (x) versus Clinical Toxicity (y). Input molecules (red stars) are shown against the open medicinal-plant reference library (USDA-Duke & MPD3 medicinal plant chemicals; n = 8272; blue). The favourable region is lower-right (high absorption, low toxicity). All panels were generated with the PangenomeAI ‘admet_screening’ pipeline (RDKit physicochemistry; ADMET-AI machine-learning endpoints; httk toxicokinetics). Raster panels are rendered at 300 dpi; vector (SVG) and an interactive HTML version of panel (D) are also provided. The reference library is an open, commercial-safe alternative to DrugBank built from medicinal-plant chemical databases.

#### 3.2.3 Cross-species ortholog mapping and molecular docking

To evaluate the evolutionary conservation of the identified compound-target interactions and support potential translational research on animal model (canine), human core targets were mapped to their respective canine orthologs. These animal-specific structures were then used to assemble a comparative docking-ready pair list. This enabled a side-by-side assessment of binding affinities and interaction fingerprints between human and dog models, providing a computational basis for cross-species efficacy and safety. Table S5 showed cross-species ortholog mapping of human and canine core targets, including the top 4 pairs by binding energy between selected phenolic compounds and core targets relevant to oxidative stress-related diseases including PTGS2, MMP2, and TNF-α. Molecular docking was used to estimate the binding affinities of three bioactive compounds toward three oxidative stress-related targets (PTGS2, MMP2, and TNF-α) in both human and canine orthologs, with more negative binding energies indicating stronger predicted interactions (Table S5). Following the widely applied interpretive convention for docking scores (with the caveat that such thresholds are software- and target-dependent; Ivanova & Karelson, 2022), values below -7.0 kcal/mol were regarded as strong, -5.0 to -7.0 kcal/mol as moderate, and above -5.0 kcal/mol as weak or potentially non-specific. The strongest interaction between a canine oxidative stress-related target and a compound was observed between PTGS2 of canine and 3,3’,4’-Tri-O-methylellagic acid at -8.641 kcal/mol. MMP2 also showed strong interactions in both human and canine orthologs, binding arjunolic acid at -6.622 kcal/mol against human ortholog and -7.667 kcal/mol against dog ortholog, and bound α-linolenic acid at -7.059 kcal/mol against the human ortholog. In contrast, TNF-α consistently yielded the weakest scores in both species (−4.349 kcal/mol for human, -3.976 kcal/mol for dog), suggesting limited draggability of this interface under the present docking conditions.

For each paired target protein, the interaction fingerprints with the phenolic compound were visualised to illustrate their binding conformations and interaction features (Figure 6). Molecular docking tested whether the compound-target relationships inferred from the network could be realised as physically plausible binding poses, focusing on the pharmacokinetically feasible pairs carried forward from the compartment screen. In the human orthologs, a strong binding observed between arjunolic acid and MMP2 (Figure 6A) was anchored by hydrogen bonds from its ring hydroxyl and carbonyl groups to Gln393 and Pro391 and by a polar contact between its carboxylate and Arg550, within a hydrophobic and van der Waals shell formed by Asn245, Phe235, Gln219 and Gly218. The second strongest pose paired 3,3’,4’-tri-O-methylellagic acid with PTGS2 (Figure 6B); the planar ellagic scaffold engaged Tyr115 and Trp100 through aromatic and hydrophobic contacts while its oxygenated substituents interacted with Arg120, residues that line the cyclo-oxygenase active site (Kurumbail et al., 1996).

**Figure 6.**
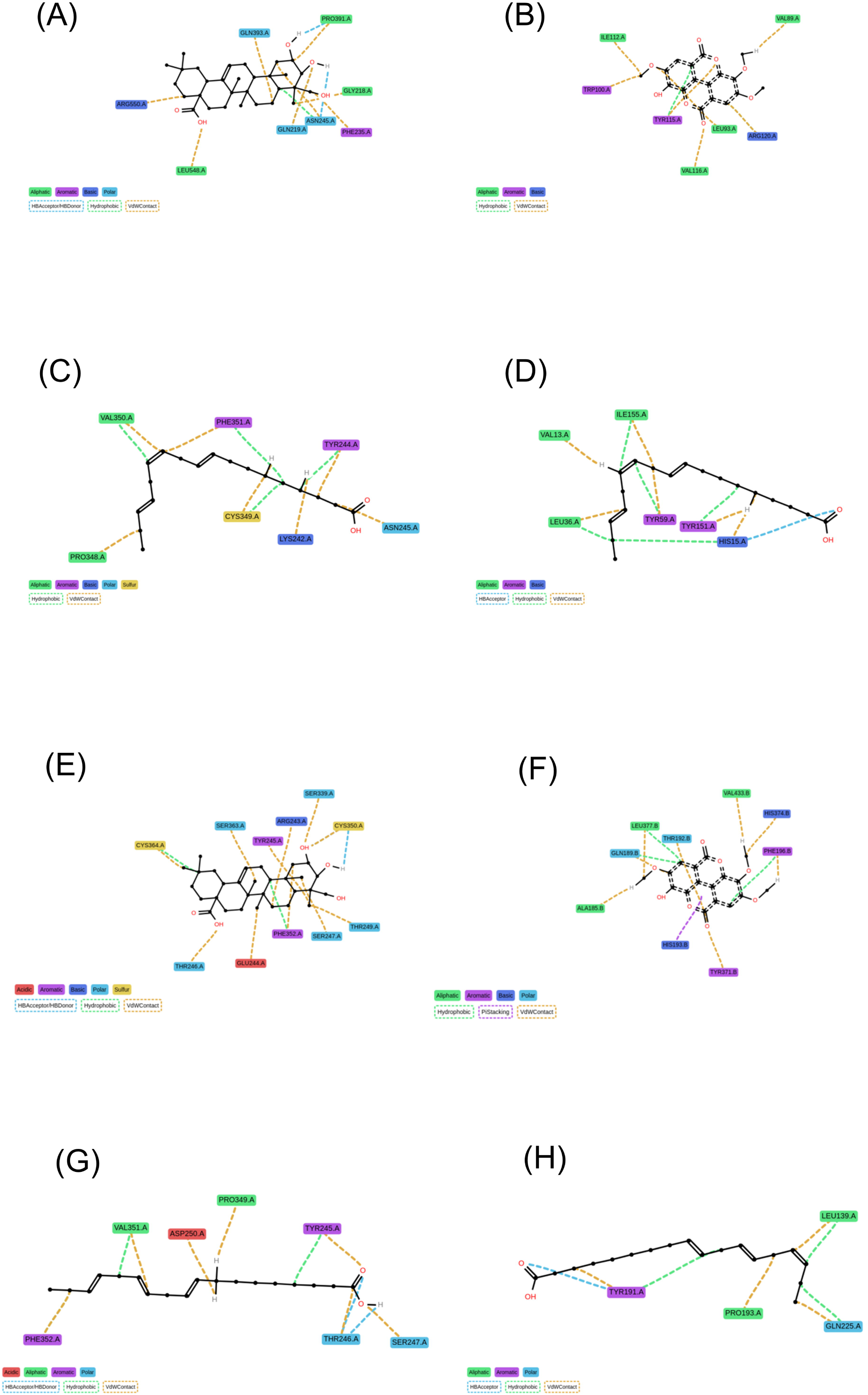
Protein-ligand interaction fingerprints for the top 4 docked compound-target pair(s). (A) MMP2 of human and Arjunolic acid (−7.06 kcal/mol). (B) PTGS2 of human and 3 3 4 - Tri-O-methylellagic acid (−6.62 kcal/mol). (C) MMP2 of human and Linolenic acid (−4.85 kcal/mol). (D) TNF of human and Linolenic acid (−4.35 kcal/mol). (E) MMP2 of dog and Arjunolic acid (−7.67 kcal/mol). (F) PTGS2 of dog and 3 3 4 -Tri-O-methylellagic acid (−8.64 kcal/mol). (G) MMP2 of dog and Linolenic acid (−4.99 kcal/mol). (H) TNF of dog and Linolenic acid (−3.98 kcal/mol). Two-dimensional interaction (LigNetwork) diagrams generated with ProLIF from the best-scoring docked pose of each pair, ordered by predicted binding affinity (most negative = strongest). Node and edge styles denote interaction types (hydrogen bonds, hydrophobic contacts, pi-stacking, cation-pi, salt bridges, halogen bonds); residue labels give the interacting protein residues.

In the canine orthologs, the same three compounds reproduced the human binding patterns with generally comparable or stronger predicted affinities (Figure 6E-6H). The strongest interaction overall was observed between the canine PTGS2 ortholog and 3,3’,4’-tri-O-methylellagic acid (−8.64 kcal/mol; Figure 6F), in which the planar ellagic scaffold occupied the cyclo-oxygenase pocket through aromatic and hydrophobic contacts analogous to those seen in the human enzyme. Arjunolic acid bound the canine MMP2 ortholog at -7.67 kcal/mol (Figure 6E), exceeding its affinity for the human ortholog and again anchored by hydrogen bonds and polar contacts from its hydroxyl, carbonyl, and carboxylate groups. In contrast, α-linolenic acid engaged canine MMP2 (−4.99 kcal/mol; Figure 6G) and TNF-alpha (−3.98 kcal/mol; Figure 6H) only weakly, consistent with the flexible acyl chain forming predominantly van der Waals contacts rather than defined hydrogen-bond networks. The close correspondence between the human and canine interaction fingerprints indicates that the key compound-target relationships are evolutionarily conserved, strengthening the translational relevance of these predictions to the dog.

The outcomes of molecular docking are corroborated by previous independent computational and mechanistic evidence for the same chemotypes. Ellagic acid and its methylated derivatives are recurrently identified as high-affinity PTGS2 ligands. Vyshnevska et al. (2022) reported that ellagic acid was docked into the COX-2 active site with a binding energy of -8.6 kcal/mol. This parallels the strong binding of 3,3’,4’-tri-O-methylellagic acid to the canine PTGS2 ortholog observed in our study (−8.64 kcal/mol; the human ortholog bound weaklier, -4.85 kcal/mol) and provides a structural rationale for the suppression of this enzymatic ROS and prostaglandin source. Asiatic acid, a close structural analogue of arjunolic acid, binds COX-2 with a docking energy of -9.80 kcal/mol and forms a stable complex over a 100-ns molecular-dynamics trajectory (Musfiroh et al., 2023). Asiatic acid derivative has been reported to down-regulate STAT3-dependent MMP-2 and MMP-9 expression (Wang et al., 2017), which mirroring the strong arjunolic acid-MMP2 interaction predicted (−7.06 kcal/mol for the human and -7.67 kcal/mol for the canine ortholog). Overall, these results indicated that 3,3’,4’-tri-O-methylellagic acid and arjunolic acid are likely the primary compounds responsible for ROS functions.

### 3.3. Intracellular antioxidant properties of Kakadu plum extract

The *in-silico* docking shows the predicted binding of chemicals with the oxidative stress-related targets. To further validate the cellular antioxidant abilities of Kakadu plum extract, *in vitro* cell studies were conducted.

#### 3.3.1. Determination of the concentration of the oxidative stress triggers H_2_O_2_

Excessive intracellular reactive oxygen species (ROS), including hydrogen peroxide (H□O□), superoxide, and hydroxyl radicals, contribute significantly to cellular injury and biological dysfunction (Sies et al., 2022). The H□O□-induced oxidative stress cell model is widely used to mimic pathological oxidative stress and cellular damage *in vitro*.

To establish an *in vitro* model of H□O□-induced intestinal oxidative stress, the cytotoxicity assay was performed on the immortalized canine small intestinal cell line to determine the H□O□ dosage. Both water and ethanol extracts of Kakadu plum were selected according to the results of the above chemical antioxidant studies. Cells were subjected to increasing oxidative stress by incubation with H□O□ at concentrations ranging from 50 µM to 1800 µM for 24 h. As shown in Figure 7A, cell viability demonstrated an overall dose-dependent decrease with increasing concentration of H□O□. A significant reduction in cell survival was first observed at 200 µM H□O□ compared to the control (p < 0.05). Cell viability of approximately 80% was observed at H□O□ concentrations of 400 - 500 µM, whilst the IC50 was observed at 900 µM H□O□. Therefore, 400 µM and 900 µM H□O□, corresponding to 80% cell viability and IC50, were selected to induce mild and severe oxidative stress injury models in the subsequent experiments.

**Figure 7.**
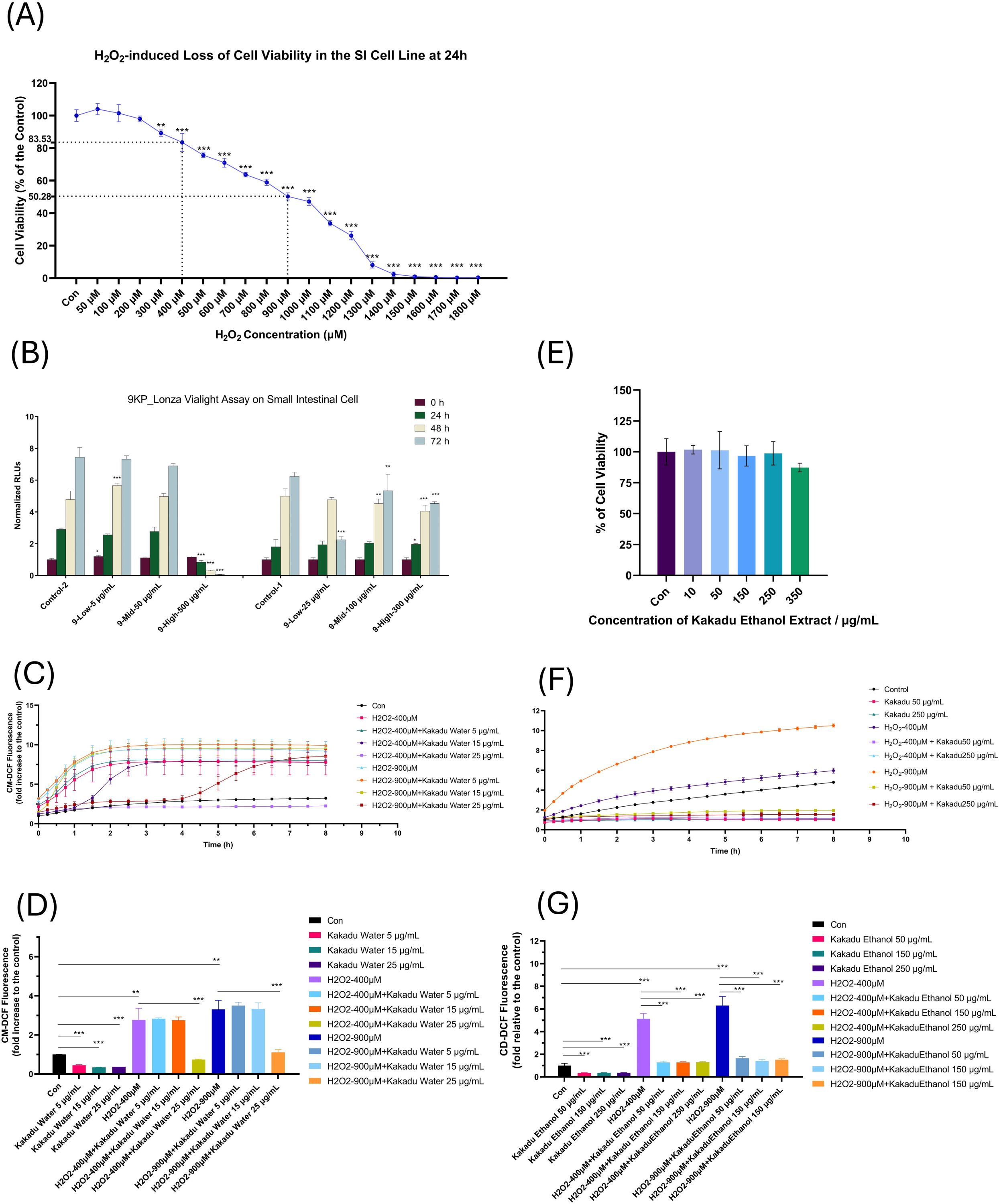
Intracellular antioxidant effect for water and ethanol extract of Kakadu plum. A: H□O□-induced loss of cell viability in the immortalized canine small intestinal epithelial cells. B: The cytotoxicity of the water extract of Kakadu plum. C: The effect of water extract of Kakadu plum on the ROS production under mild and severe oxidative stress model. D: The inhibitory effect of water extract of Kakadu plum on the ROS production under oxidative stress for 4h. E: The cytotoxicity of the ethanol extract of Kakadu plum. F: The effect of ethanol extract of Kakadu plum on the ROS production under mild and severe oxidative stress model. G: The inhibitory effect of ethanol extract of Kakadu plum on the ROS production under oxidative stress for 4h. Cells were seeded at densities of 2.0×104 in 96-well plates and incubated for 24 h with increasing concentrations of H□O□. The data are normalized as the relative response of treated cells as compared to the untreated control from triplicate experiments. Statistical significance is given as follows: * p < 0.05, ** p < 0.001, and *** p < 0.005, as compared to untreated controls.

The sensitivity and tolerance of intestinal epithelial cells to H□O□-induced oxidative stress vary considerably across cell lines of different origins and species. Previous research by Mohammed et al. (2023) demonstrated that immortalized normal oral cells were more sensitive to H□O□ than malignant oral epithelial cells. The IC50 value of 800 µM was observed in the two malignant cell lines H400 and H357, which was two-fold that of the immortalized normal oral cell line OKF6 (400 µM), indicating greater resistance to H□O□-induced cytotoxicity in malignant cells. Among human malignant intestinal cell lines, Caco-2 cells (human colorectal adenocarcinoma) have been widely employed in H□O□-induced oxidative stress models. C. Li et al. (2023) determined an IC50 of approximately 800 µM and 80% cell viability at 600 µM H□O□ using the MTT assay, whilst Su et al. (2025) reported 56.52% cell viability at 800 µM H□O□ using the CCK-8 assay, with an IC50 of approximately 900 µM. For the HT-29 cell line (human colorectal adenocarcinoma), (Park et al., 2025) observed an IC50 of 1000 µM H□O□ and 80% cell viability at around 1250 µM via the MTT assay, suggesting greater H□O□ tolerance compared with Caco-2 cells. Studies using normal human intestinal epithelial cell lines are comparatively limited. The NCM460 cell line, an immortalized cell line derived from normal human colon mucosal epithelial cells, exhibited a reduction in cell viability to 75.16% when treated with 1000 µM H□O□ for 4 h (Yao et al., 2026). There are other H□O□-induced oxidative stress cell models using normal intestinal epithelial cell lines from other animal species. For the normal porcine intestinal epithelial cell line IPEC-J2, Gao et al. (2026) determined an IC50 of 400-600 µM H□O□ using the CCK-8 assay, whilst Wang et al. (2021) obtained a higher IC50 of 1000 µM using the MTT assay. For immortalized normal bovine intestinal epithelial cells, an IC50 of 1000 µM H□O□ using the CCK-8 assay was determined and employed in the oxidative apoptosis model (Mao et al., 2022). In contrast, ovine intestinal epithelial cells demonstrated considerably higher sensitivity to H□O□, with an IC50 of 150 µM H□O□ reported by Zhang et al. (2021). For the immortalized normal rat small intestinal epithelial cell line IEC-6, H□O□ concentrations of 300-600 µM were required to reduce cell viability to approximately 70%, with Peng et al. (2022) reporting 70% cell viability at 300 µM H□O□, whilst Jia et al. (2021) reported approximately 70% viability at 600 µM H□O□ and 62% viability at 1000 µM H□O□. In the present study, the immortalized canine small intestinal cell line exhibited an IC50 of 900 µM H□O□ and 80% cell viability at 400 µM H□O□, values consistent with those reported for human malignant intestinal cell lines (Caco-2: IC50 at 900 µM H□O□; HT-29: IC50 at 1000 µM) and bovine intestinal epithelial cells (IC50: 1000 µM H□O□), suggesting a comparatively higher tolerance to H□O□-induced oxidative stress than normal intestinal epithelial cell lines of other species such as ovine (IC50 at 150 µM H□O□) and porcine (IC50: 400-600 µM H□O□) cells.

#### 3.3.2 Cytotoxicity and kinetics of cellular ROS of Kakadu plum water and ethanol extract

To determine the cytotoxic potential of the Kakadu plum extract, Lonza ViaLight® Plus cell proliferation cytotoxicity assay was performed on the canine immortalized small intestinal cell line. For the water extract, cells were treated with concentrations ranging from 5 to 500 µg/mL across four timepoints (0, 24, 48, and 72 h). As shown in Figure 7B, concentrations lower than 50 µg/mL maintained normal cell growth and proliferation patterns; cell viability was maintained above 80% at doses below 100 µg/mL, whereas a significant dose-dependent reduction in cell viability was observed when concentrations exceeded 100 µg/mL. Therefore, doses below 50 µg/mL were selected for subsequent experiments.

Both 400 µM and 900 µM H□O□ were employed as mild and severe oxidative stress models, respectively, with ROS production and real-time kinetics evaluated using the CM-H2DCFDA fluorescent probe (Figure 7). The effect of Kakadu plum water extract (5-25 µg/mL) on H□O□-induced ROS production over 8 h is presented in Figure 7C and Figures S4A-S4C. Under both models, ROS increased progressively, peaking at 3.5 h for 900 µM H□O□ (approximately 9.4-fold of the control at time 0) and at 4 h for the 400 µM H□O□ (approximately 7.9-fold of the control at time 0) before stabilizing. At 4 h, cellular ROS production under 900 µM H□O□ was 1.2-fold that under 400 µM H□O□. Under mild oxidative stress (400 µM H□O□), water extract of Kakadu plum at 15 and 25 µg/mL strongly inhibited ROS production throughout. Interestingly, at 5 µg/mL, ROS was initially suppressed but rose from 1 h to 3 h until reaching levels comparable to the 400 µM H□O□ control. Under severe oxidative stress (900 µM H□O□), only water extract of Kakadu plum at 25 µg/mL sustained ROS inhibition over 8 h, which reduced H□O□-induced ROS 3.3-fold at 2 h and 3.0-fold at 4 h. At 5 µg/mL, no inhibitory effect was observed, whilst 15 µg/mL suppressed ROS only within the first 4 h, after which ROS increased until 8 h. Notably, in the absence of oxidative stress, all tested concentrations (5-25 µg/mL) markedly reduced intracellular ROS levels (Figure S4C). The inhibitory effects of Kakadu plum water extract at concentrations of 5 to 25 µg/mL on ROS production at 2 h and 4 h are presented in Figure S4D and Figure 7D. The Kakadu plum water extract at low concentrations (both 5 and 15 µg/mL) showed no significant inhibitory effect on ROS at 2 h or 4 h. Only the 25 µg/mL suppressed ROS production to 0.28-fold of the 400 µM H□O□ control at 2 h, and to 0.31-fold of the 900 µM H□O□ control at 2 h (Figure S4D). The water extract at 25 µg/mL suppressed ROS production to 0.27-fold of the 400 µM H□O□ control and 0.33-fold of the 900 µM H₂O₂ control at 4 h, respectively (Figure 7D).

The cell cytotoxicity assay of the Kakadu plum ethanol extract at 10 to 350 µg/mL was assessed after 24 h of treatment (Figure 7E). Compared with the water extract, cells exhibited greater tolerance to the ethanol extract, with a noticeable reduction in cell viability observed only at concentrations exceeding 250 µg/mL. Consequently, doses below 250 µg/mL were selected for subsequent experiments to maintain cell viability above 80% and to exclude confounding cytotoxic effects. The inhibitory effect of Kakadu plum ethanol extract (50-250 µg/mL) on H□O□-induced ROS production (400 µM and 900 µM H□O□) using the CM-H2DCFDA fluorescent probe over 8 h is presented in Figure 7F and Figures S4E-S4G. The ROS production of 900 µM H□O□ peaked at 3.5 h, which was 2.0-fold that of the 400 µM H□O□ and 8.4-fold of the control. In the absence of oxidative stress, both tested concentrations (50 and 250 µg/mL) markedly reduced intracellular ROS levels (Figure S4G). Under both the mild oxidative stress (400 µM H□O□) and severe oxidative stress (900 µM H□O□), ethanol extract of Kakadu plum at both 50 and 250 µg/mL strongly inhibited ROS production throughout. The dose-dependent inhibitory effects were observed at concentrations of 50 to 250 µg/mL. At 2h (Figure S4H), the ethanol extract at 50 and 250 µg/mL suppressed ROS production to 0.35-fold and 0.31-fold of the 400 µM H□O□ control, respectively, and to 0.23-fold and 0.21-fold under 900 µM H□O□. Similarly, at 4h (Figure 7G), the ethanol extract at 50 and 250 µg/mL inhibited ROS production to 0.25-fold and 0.25-fold of the 400 µM H□O□ control, respectively, and to 0.20-fold and 0.17-fold under 900 µM H□O□.

The Kakadu plum extracts were well tolerated by the immortalised canine small intestinal cells, with the ethanol extract permitting substantially higher exposure (≤250 µg/mL) than the water extract (≤50 µg/mL). These thresholds are consistent with the wide range of cytotoxic concentrations reported for Kakadu plum across cell types and preparations. Ramadhania et al. (2022) found that a water extract significantly reduced the viability of RAW 264.7 murine macrophages and A549 human lung carcinoma cells at 10 µg/mL as assessed by MTT assay, whereas the immortalized normal human breast epithelial line MCF-10A tolerated up to approximately 630 µg/mL, with a notably high half-maximal cytotoxic concentration (CC□□) near 5,000 µg/mL (Alwis et al., 2025). The extraction solvent has also been reported to have a significant impact on cytotoxicity; the aqueous acidified ethanol extract of Kakadu plum (CC□□ = 1676-2456 μg/mL) exhibited much lower CCCC values against HepG-2 cells than the water extract (CC□□ = 5440-7337 μg/mL) (Bobasa et al., 2022).

The suppression of intracellular ROS by Kakadu plum extracts is consistent with the cytoprotective effects reported for other polyphenol-rich plant extracts in H□O□-triggered intestinal epithelial models. In the porcine intestinal epithelial cell line IPEC-J2, green tea polyphenols (100 µg/mL) decreased intracellular ROS in epithelial cells exposed to 500 µM H□O□ by engaging the ERK1/2-NFE2L2-HMOX1 axis (Ma et al., 2021), whereas tea polyphenols (200 mg/L) restored redox balance in fluoride-stressed IPEC-J2 cells (Xie et al., 2025). In the human colonic cancer cell Caco-2 model, proanthocyanidin-rich grape seed extract reduced ROS production and restored tight-junction barrier function under inflammatory-oxidative challenge (Nallathambi et al., 2020). Blueberry anthocyanins similarly lowered ROS and malondialdehyde and up-regulated superoxide dismutase, catalase, and glutathione peroxidase in H□OC-injured epithelial cells (Huang et al., 2018). The composition of the extracts supports this interpretation. Hydrolysable tannins such as corilagin, chebulagic acid, and chebulinic acid, together with ellagic acid derivatives and triterpenoids, are prominent constituents of Kakadu plum (Adiamo et al., 2024), and the network pharmacology analysis indicated that these phenolics converge on a coherent inflammation-redox module (IL6, PTGS2, STAT3, TNF, AKT1, SRC, HSP90AA1, MMP2, and NOS2) rather than acting solely as radical scavengers. Molecular docking further predicted that key constituents 3,3’,4’-tri-O-methylellagic acid and arjunolic acid, bind conserved canine orthologs of PTGS2 and MMP2 with moderate-to-strong affinity, providing a plausible structural basis for the suppression of enzymatic ROS sources and inflammatory signalling in addition to direct radical scavenging. According to the predicted chemical-target interaction, both 3,3’,4’-tri-O-methylellagic acid and arjunolic acid have documented cellular antioxidant mechanisms that plausibly underlie the ROS suppression observed here. The 3,3’,4’-tri-O-methylellagic acid is a permethylated derivative of ellagic acid, and the parent scaffold is an established cytoprotectant that acts on two levels: it directly scavenges hydroxyl, peroxyl, and superoxide radicals and inhibits lipid peroxidation, and at the intracellular level, it activates the Keap1-Nrf2/ARE axis to induce endogenous antioxidant enzymes (e.g., HO-1, NQO1) while suppressing IκBα/NF-κB signalling and the downstream cytokines TNF-α and IL-6 (Ebrahimi et al., 2019; Wojtunik-Kulesza et al., 2025). Notably, ellagic acid itself is poorly soluble and poorly absorbed (aqueous solubility ≈ 9.7 µg/mL; oral bioavailability < 1%), whereas O-methylation of its phenolic hydroxyls increases lipophilicity and membrane permeability; the methylated derivative would therefore be expected to partition more readily into the epithelial cytosol and reach intracellular enzymatic targets such as PTGS2, consistent with its strong predicted binding and with the IL6-PTGS2-STAT3-TNF module recovered by the network analysis. Arjunolic acid, contributes a complementary mechanism: it quenches superoxide, hydroxyl, H□O□, and nitric oxide radicals, chelates redox-active transition metals through its vicinal hydroxyl groups (limiting Fenton-type ·OH generation), and preserves mitochondrial membrane potential to block mitochondria-mediated, caspase-dependent apoptosis (Ghosh & Sil, 2013). Thus, within this predicted cytoprotective repertoire, including direct radical scavenging, Nrf2-driven enzyme induction, and mitochondrial preservation, the direct antioxidant mechanisms likely dominate the intracellular ROS attenuation, while the enzymatic-target modulation inferred from the *in silico* arjunolic acid-MMP2 and methylellagic acid-PTGS2 interactions represents a plausible signalling-level contribution that remains to be validated experimentally. These results indicated Kakadu plum extract as a multi-target antioxidant capable of attenuating epithelial oxidative stress.

## 4. Conclusions

Kakadu plum extracts demonstrated suppressing effects on intracellular oxidative stress in the canine intestinal epithelial cell model. Extraction solvent (hydroalcoholic solvents such as ethanol and methanol), instead of processing technique, was the principal determinant of antioxidant capacity. By integrating chemical profiling, network pharmacology, ADMET profiling, cross-species ortholog mapping, and molecular docking, the identified phenolics were linked to ROS related targets, and the binding of key constituents to canine orthologs of PTGS2 and MMP2 were confirmed *in silico*. In the H□O□-induced oxidative-stress cell model, both the water and ethanol extracts of Kakadu plum significantly suppressed intracellular ROS in a dose dependent manner, with the ethanol extract effective across a wider concentration range. The chemical, computational, and cellular findings provide a mechanistic and translational rationale for developing Kakadu plum as a natural, multi-target antioxidant ingredient for canine intestinal health. These conclusions are based on commercial Kakadu plum powder, the phenolic profile was characterised qualitatively without individual quantification due to lack of individual standards, and the network-pharmacology and docking predictions were not experimentally validated; these limitations should be addressed in future work. Future research would aim at experimentally validating the predicted molecular interactions, elucidating the underlying molecular mechanisms through whole-transcriptome analysis, and evaluating the effects on intestinal barrier function, inflammatory responses, and *in vivo* efficacy in animal models.

## Acknowledgements

We acknowledge the Traditional Owners of Australia and their continuing connection to the land, waters, and community. We also sincerely thank all the researchers and technicians from Melbourne Dental School, Rita Paolini, Syed Ameer Hamza, Caroline Moore, Sze Wei Liu, Su Toulson for all the support. We thank the Walter and Eliza Hall Institute for kindly providing the SV-40 virus. We thank Cai Shen and Ling-Zhi Cheong for the assistance of FTIR analysis.

This work was supported by a Melbourne Research Scholarship, together with funding from Australia Talentail Pty Ltd as part of an ongoing research partnership with the University of Melbourne (Project Cayuse ID: <u>25-7791 and 26-2438</u>). In particular, we thank Ms Yolanda Franklyn for her valuable industry guidance and continued support throughout this project. AI agent skills (academic-skills-food-nutrition, v1.30.0) developed by our research group was used on Anthropic claude desktop using optus 4.8 base model to support manuscript reviewing and editing process (Zhang et al., 2026). All information suggested by AI has been reviewed by authors to validate the accuracy.

## CRediT authorship contribution statement

**Yidan He**: Conceptualization, Methodology, Formal analysis, Data curation, Validation, Writing - original draft. **Xuefu Zhou**: Software, Methodology, Validation, Writing - review & editing. **Antonio Celentano**: Conceptualization, Supervision, Methodology, Resources, Writing - review & editing. **Nicola Cirillo**: Conceptualization, Methodology, Supervision, Writing - review & editing. **Long Cheng**: Conceptualization, Methodology, Supervision, Writing - review & editing. **Zhongxiang Fang**: Conceptualization, Methodology, Supervision, Writing - review & editing. **Pangzhen Zhang**: Conceptualization, Methodology, Software, Resources, Writing - review & editing, Supervision, Project administration, Funding acquisition

## Data availability

The data supporting the findings of this study are available from the corresponding author upon reasonable request. The information about the *in silico* analysis pipeline will be made available to the public on the PangenomeAI platform (https://github.com/PangenomeAI).

## Supplementary data

## References

1. Adiamo, O. Q., Bobasa, E. M., Phan, A. D. T., Akter, S., Seididamyeh, M., Dayananda, B., Gaisawat, M. B., Kubow, S., Sivakumar, D., & Sultanbawa, Y. (2024). In-vitro colonic fermentation of Kakadu plum (Terminalia ferdinandiana) fruit powder: Microbial biotransformation of phenolic compounds and cytotoxicity. Food Chem, 448, 139057. 10.1016/j.foodchem.2024.139057

2. Akter, R., Kwak, G.-Y., Ahn, J. C., Mathiyalagan, R., Ramadhania, Z. M., Yang, D. C., & Kang, S. C. (2021). Protective Effect and Potential Antioxidant Role of Kakadu Plum Extracts on Alcohol-Induced Oxidative Damage in HepG2 Cells. Applied Sciences, 12(1). 10.3390/app12010236

3. Akter, S., Netzel, M. E., Fletcher, M. T., Tinggi, U., & Sultanbawa, Y. (2018). Chemical and Nutritional Composition of Terminalia ferdinandiana (Kakadu Plum) Kernels: A Novel Nutrition Source. Foods, 7(4). 10.3390/foods7040060

4. Akter, S., Netzel, M. E., Tinggi, U., Osborne, S. A., Fletcher, M. T., & Sultanbawa, Y. (2019). Antioxidant Rich Extracts of Terminalia ferdinandiana Inhibit the Growth of Foodborne Bacteria. Foods, 8(8). 10.3390/foods8080281

5. Alwis, W. H. S., Murthy, V., Wang, H., Khandanlou, R., & Weir, R. (2025). Biofabrication of Terminalia ferdinandiana-Conjugated Gold Nanoparticles and Their Anticancer Properties. Life (Basel), 15(12). 10.3390/life15121829

6. Amiri, M., Arab, M., Khalili Sadrabad, E., Mollakhalili-Meybodi, N., & Fallahzadeh, H. (2023). Effect of gamma irradiation treatment on the antioxidant activity, phenolic compounds and flavonoid content of common buckwheat. Radiation Physics and Chemistry, 212. 10.1016/j.radphyschem.2023.111127

7. Bobasa, E. M., Akter, S., Phan, A. D. T., Netzel, M. E., Cozzolino, D., Osborne, S., & Sultanbawa, Y. (2022). Impact of Growing Location on Kakadu Plum Fruit Composition and In Vitro Bioactivity as Determinants of Its Nutraceutical Potential. Nutraceuticals, 3(1), 13–25. 10.3390/nutraceuticals3010002

8. Chaliha, M. (2018). Exploring the bioactive potential of Terminalia ferdinandiana (Kakadu Plum)-a native plant of Australia. 10.14264/6663937

9. Chen, J., Yang, J., Ma, L., Li, J., Shahzad, N., & Kim, C. K. (2020). Structure-antioxidant activity relationship of methoxy, phenolic hydroxyl, and carboxylic acid groups of phenolic acids. Sci Rep, 10(1), 2611. 10.1038/s41598-020-59451-z

10. Cock, I. E. (2015). The medicinal properties and phytochemistry of plants of the genus Terminalia (Combretaceae). Inflammopharmacology, 23(5), 203–229. 10.1007/s10787-015-0246-z

11. Corcoran, A., & Cotter, T. G. (2013). Redox regulation of protein kinases. FEBS J, 280(9), 1944–1965. 10.1111/febs.12224

12. Ebrahimi, R., Sepand, M. R., Seyednejad, S. A., Omidi, A., Akbariani, M., Gholami, M., & Sabzevari, O. (2019). Ellagic acid reduces methotrexate-induced apoptosis and mitochondrial dysfunction via up-regulating Nrf2 expression and inhibiting the IkBalpha/NFkB in rats. Daru, 27(2), 721–733. 10.1007/s40199-019-00309-9

13. Gao, M., Xie, Z., Mao, P., Zhang, X., Zhao, L., & Ma, W. (2026). The protective effects and mechanism of N-carbamoyl glutamate against H(2)O(2)-induced oxidative damage in IPEC-J2 cell. BMC Vet Res, 22(1). 10.1186/s12917-025-05272-z

14. Ghosh, J., & Sil, P. C. (2013). Arjunolic acid: a new multifunctional therapeutic promise of alternative medicine. Biochimie, 95(6), 1098–1109. 10.1016/j.biochi.2013.01.016

15. He, Y., Fang, Z., Ying, D., Franklyn, Y., & Zhang, P. (2023). Terminalia ferdinandiana Exell (Kakadu plum): Nutritional value, phenolic compounds, health benefits and potential industrial applications. Food Bioscience, 56. 10.1016/j.fbio.2023.103427

16. Hernández-Corroto, E., Boussetta, N., Marina, M. L., García, M. C., & Vorobiev, E. (2022). High voltage electrical discharges followed by deep eutectic solvents extraction for the valorization of pomegranate seeds (Punica granatum L.). Innovative Food Science & Emerging Technologies, 79. 10.1016/j.ifset.2022.103055

17. Hopkins, A. L. (2008). Network pharmacology: the next paradigm in drug discovery. Nat Chem Biol, 4(11), 682–690. 10.1038/nchembio.118

18. Hossain, M. A., Arafat, M. Y., Alam, M., & Hossain, M. M. (2021). Effect of solvent types on the antioxidant activity and total flavonoids of some Bangladeshi legumes. Food Research, 5(4), 329–335. 10.26656/fr.2017.5(4).035

19. Huang, W. Y., Wu, H., Li, D. J., Song, J. F., Xiao, Y. D., Liu, C. Q., Zhou, J. Z., & Sui, Z. Q. (2018). Protective Effects of Blueberry Anthocyanins against H(2)O(2)-Induced Oxidative Injuries in Human Retinal Pigment Epithelial Cells. J Agric Food Chem, 66(7), 1638–1648. 10.1021/acs.jafc.7b06135

20. Jia, Y., Wang, Y., Li, R., Li, S., Zhang, M., He, C., & Chen, H. (2021). The structural characteristic of acidic-hydrolyzed corn silk polysaccharides and its protection on the H(2)O(2)-injured intestinal epithelial cells. Food Chem, 356, 129691. 10.1016/j.foodchem.2021.129691

21. Khaligh, S. F., & Asoodeh, A. (2022). Green synthesis and biological characterization of cerium oxide nanoemulsion against human HT-29 colon cancer cell line. Materials Technology, 37(12), 2318–2338. 10.1080/10667857.2022.2031492

22. Konczak, I., Maillot, F., & Dalar, A. (2014). Phytochemical divergence in 45 accessions of Terminalia ferdinandiana (Kakadu plum). Food Chem, 151, 248–256. 10.1016/j.foodchem.2013.11.049

23. Konczak, I., Zabaras, D., Dunstan, M., & Aguas, P. (2010). Antioxidant capacity and hydrophilic phytochemicals in commercially grown native Australian fruits. Food Chemistry, 123(4), 1048–1054. 10.1016/j.foodchem.2010.05.060

24. Kumar, P. S., Kodi, T., Gopinathan, A., Byregowda, B. H., Nandakumar, K., & Kishore, A. (2026). New Insights on Heat Shock Proteins as Regulators of Reactive Oxygen Species Across Various Stressors in Diseases. Cell Biochem Funct, 44(2), e70173. 10.1002/cbf.70173

25. Kurumbail, R. G., Stevens, A. M., Gierse, J. K., McDonald, J. J., Stegeman, R. A., Pak, J. Y., Gildehaus, D., iyashiro, J. M., Penning, T. D., Seibert, K., Isakson, P. C., & Stallings, W. C. (1996). Structural basis for selective inhibition of cyclooxygenase-2 by anti-inflammatory agents. Nature, 384(6610), 644–648. 10.1038/384644a0

26. Landete, J. M. (2011). Ellagitannins, ellagic acid and their derived metabolites: A review about source, metabolism, functions and health. Food Research International, 44(5), 1150–1160. 10.1016/j.foodres.2011.04.027

27. Lang, Y., Gao, N., Zang, Z., Meng, X., Lin, Y., Yang, S., Yang, Y., Jin, Z., & Li, B. (2024). Classification and antioxidant assays of polyphenols: a review. Journal of Future Foods, 4(3), 193–204. 10.1016/j.jfutfo.2023.07.002

28. Leonard, W., Zhang, P., Ying, D., Xiong, Y., & Fang, Z. (2021). Extrusion improves the phenolic profile and biological activities of hempseed (Cannabis sativa L.) hull. Food Chem, 346, 128606. 10.1016/j.foodchem.2020.128606

29. Li, C., Chen, X., Li, L., Cheng, J., Chen, H., Gao, Q., Yang, F., Cai, X., & Wang, S. (2023). Protective effect of antioxidant peptides from bass (Lateolabrax japonicus) on oxidative stress injury in CacoC2 cells. Food Frontiers, 4(2), 818–830. 10.1002/fft2.224

30. Li, J., Liu, H., Mazhar, M. S., Quddus, S., & Suleria, H. A. R. (2023). LC-ESI-QTOF-MS/MS profiling of phenolic compounds in Australian native plums and their potential antioxidant activities. Food Bioscience, 52. 10.1016/j.fbio.2022.102331

31. Ma, Y., Ma, X., An, Y., Sun, Y., Dou, W., Li, M., Bao, H., & Zhang, C. (2021). Green Tea Polyphenols Alleviate Hydrogen Peroxide-Induced Oxidative Stress, Inflammation, and Apoptosis in Bovine Mammary Epithelial Cells by Activating ERK1/2-NFE2L2-HMOX1 Pathways. Front Vet Sci, 8, 804241. 10.3389/fvets.2021.804241

32. Makanjuola, S. A. (2017). Influence of particle size and extraction solvent on antioxidant properties of extracts of tea, ginger, and tea-ginger blend. Food Sci Nutr, 5(6), 1179–1185. 10.1002/fsn3.509

33. Mao, H., Zhang, Y., Ji, W., Yun, Y., Wei, X., Cui, Y., & Wang, C. (2022). Leucine protects bovine intestinal epithelial cells from hydrogen peroxide-induced apoptosis by alleviating oxidative damage. J Sci Food Agric, 102(13), 5903–5912. 10.1002/jsfa.11941

34. Merecz-Sadowska, A., Sadowski, A., Zielinska-Blizniewska, H., Zajdel, K., & Zajdel, R. (2025). Network Pharmacology as a Tool to Investigate the Antioxidant and Anti-Inflammatory Potential of Plant Secondary Metabolites-A Review and Perspectives. Int J Mol Sci, 26(14). 10.3390/ijms26146678

35. Mohammed, A. I., Sangha, S., Nguyen, H., Shin, D. H., Pan, M., Park, H., McCullough, M. J., Celentano, A., & Cirillo, N. (2023). Assessment of Oxidative Stress-Induced Oral Epithelial Toxicity. Biomolecules, 13(8). 10.3390/biom13081239

36. Musfiroh, I., Kartasasmita, R. E., Ibrahim, S., Muchtaridi, M., Hidayat, S., & Ikram, N. K. K. (2023). Stability Analysis of the Asiatic Acid-COX-2 Complex Using 100 ns Molecular Dynamic Simulations and Its Selectivity against COX-2 as a Potential Anti-Inflammatory Candidate. Molecules, 28(9). 10.3390/molecules28093762

37. Nallathambi, R., Poulev, A., Zuk, J. B., & Raskin, I. (2020). Proanthocyanidin-Rich Grape Seed Extract Reduces Inflammation and Oxidative Stress and Restores Tight Junction Barrier Function in Caco-2 Colon Cells. Nutrients, 12(6). 10.3390/nu12061623

38. Park, K. M., Lee, Y. R., Lee, N. K., & Paik, H. D. (2025). Lactiplantibacillus plantarum WB3801 and Lactiplantibacillus plantarum WB3808 Showed Antioxidant Effect and Anti-Apoptosis through Activation of the Keap1/Nrf2/HO-1 Pathway in H(2)O(2)-Induced HT-29 Cells. J Microbiol Biotechnol, 35, e2508014. 10.4014/jmb.2508.08014

39. Peng, B., Cai, B., & Pan, J. (2022). Octopus-derived antioxidant peptide protects against hydrogen peroxide-induced oxidative stress in IEC-6 cells. Food Sci Nutr, 10(11), 4049–4058. 10.1002/fsn3.3000

40. Pfundstein, B., El Desouky, S. K., Hull, W. E., Haubner, R., Erben, G., & Owen, R. W. (2010). Polyphenolic compounds in the fruits of Egyptian medicinal plants (Terminalia bellerica, Terminalia chebula and Terminalia horrida): characterization, quantitation and determination of antioxidant capacities. Phytochemistry, 71(10), 1132–1148. 10.1016/j.phytochem.2010.03.018

41. Phan, A. D. T., Zhang, J., Seididamyeh, M., Srivarathan, S., Netzel, M. E., Sivakumar, D., & Sultanbawa, Y. (2022). Hydrolysable tannins, physicochemical properties, and antioxidant property of wild-harvested Terminalia ferdinandiana (exell) fruit at different maturity stages. Front Nutr, 9, 961679. 10.3389/fnut.2022.961679

42. Ramadhania, Z. M., Nahar, J., Ahn, J. C., Yang, D. U., Kim, J. H., Lee, D. W., Kong, B. M., Mathiyalagan, R., Rupa, E. J., Akter, R., Yang, D. C., Kang, S. C., & Kwak, G.-Y. (2022). Terminalia ferdinandiana (Kakadu Plum)-Mediated Bio-Synthesized ZnO Nanoparticles for Enhancement of Anti-Lung Cancer and Anti-Inflammatory Activities. Applied Sciences, 12(6). 10.3390/app12063081

43. Reuter, S., Gupta, S. C., Chaturvedi, M. M., & Aggarwal, B. B. (2010). Oxidative stress, inflammation, and cancer: how are they linked? Free Radic Biol Med, 49(11), 1603–1616. 10.1016/j.freeradbiomed.2010.09.006

44. Sies, H., Belousov, V. V., Chandel, N. S., Davies, M. J., Jones, D. P., Mann, G. E., Murphy, M. P., Yamamoto, M., & Winterbourn, C. (2022). Defining roles of specific reactive oxygen species (ROS) in cell biology and physiology. Nat Rev Mol Cell Biol, 23(7), 499–515. 10.1038/s41580-022-00456-z

45. Su, J., Lu, J., Zeng, H., Sun, W., Zhu, X., Yang, R., Qu, L., & Zhao, C. (2025). Identification and in silico analysis of novel antioxidant peptides from Grifola frondosa hydrolysates: Cytoprotective effects in H(2)O(2)-induced Caco-2 cells. Food Res Int, 221(Pt 1), 117273. 10.1016/j.foodres.2025.117273

46. Theocharis, G., & Andlauer, W. (2013). Innovative microwave-assisted hydrolysis of ellagitannins and quantification as ellagic acid equivalents. Food Chem, 138(4), 2430–2434. 10.1016/j.foodchem.2012.12.015

47. Vyshnevska, L., Severina, H. I., Prokopenko, Y., & Shmalko, A. (2022). Molecular docking investigation of anti-inflammatory herbal compounds as potential LOX-5 and COX-2 inhibitors. Pharmacia, 69(3), 733–744. 10.3897/pharmacia.69.e89400

48. Wang, G., Jing, Y., Cao, L., Gong, C., Gong, Z., & Cao, X. (2017). A novel synthetic Asiatic acid derivative induces apoptosis and inhibits proliferation and mobility of gastric cancer cells by suppressing STAT3 signaling pathway. Onco Targets Ther, 10, 55–66. 10.2147/OTT.S121619

49. Wang, J., Zhang, W., Wang, S., Wang, Y., Chu, X., & Ji, H. (2021). Lactobacillus plantarum Exhibits Antioxidant and Cytoprotective Activities in Porcine Intestinal Epithelial Cells Exposed to Hydrogen Peroxide. Oxid Med Cell Longev, 2021, 8936907. 10.1155/2021/8936907

50. Wojtunik-Kulesza, K., Nizinski, P., Krajewska, A., Oniszczuk, T., Combrzynski, M., & Oniszczuk, A. (2025). Therapeutic Potential of Ellagic Acid in Liver Diseases. Molecules, 30(12). 10.3390/molecules30122596

51. Xie, C., Niu, S., & Tian, W. (2025). Tea Polyphenols Relieve the Fluoride-Induced Oxidative Stress in the Intestinal Porcine Epithelial Cell Model. Toxics, 13(2). 10.3390/toxics13020083

52. Yao, Y., Wen, X., He, X., Liao, D., Li, M., Fan, J., Liang, R., Huang, X., & Li, N. (2026). Protective Effect of Gastrodia elata Polysaccharide GEP-2 Against Oxidative Stress in Intestinal Epithelial NCM460 Cells. Int J Mol Sci, 27(6). 10.3390/ijms27062655

53. Zhang, H., Zhang, Y., Liu, X., Elsabagh, M., Yu, Y., Peng, A., Dai, S., & Wang, H. (2021). L-Arginine inhibits hydrogen peroxide-induced oxidative damage and inflammatory response by regulating antioxidant capacity in ovine intestinal epithelial cells. Italian Journal of Animal Science, 20(1), 1620–1632. 10.1080/1828051x.2021.1973916

54. Zhang, P., Liang, Z., Huang, P., Shen, C., & Yao, X. (2026). Academic Skills for Food & Nutrition Science (Version 1.35.0) [Computer software]. 10.5281/zenodo.21372994

