## Supplementary tables for "Optimized Kakadu Plum Extracts Inhibit Intracellular Oxidative Stress in Canine Small Intestinal Cell Model"

Table S1. The antioxidant properties (i.e., TPC, TFC, DPPH, and FRAP) for free, bound, and total phenolics of Kakadu extracts using different solvents.

|  | Variable | Ethanol | Methanol | Water | p-value |
| --- | --- | --- | --- | --- | --- |
| Free | TPC_Free (mg GAE/g DW) | 225.25 Â± 14.18 a | 204.75 Â± 10.52 b | 205.68 Â± 6.56 b | 0.0006238 *** |
|  | TFC_Free (mg CAE/g DW) | 6.07 Â± 0.60 a | 5.93 Â± 1.32 a | 2.59 Â± 0.15 b | 3.021e-09 *** |
|  | DPPH_Free (mg Trolox eq./g DW) | 259.22 Â± 22.55 a | 264.13 Â± 20.25 a | 181.36 Â± 36.64 b | 1.063e-06 *** |
|  | FRAP_Free µmol Fe2+ eq./g DW) | 1625.98 Â± 179.46 b | 1770.10 Â± 206.07 b | 2201.99 Â± 118.74 a | 6.513e-07 *** |
| Bound | TPC_Bound (mg GAE/g DW) | 4.78 Â± 1.09 b | 5.21 Â± 0.90 b | 8.26 Â± 0.82 a | 0.0001171 *** |
|  | TFC_Bound (mg CAE/g DW) | 0.31 Â± 0.14 b | 0.23 Â± 0.08 b | 0.65 Â± 0.15 a | 4.942e-07 *** |
|  | DPPH_Bound (mg Trolox eq./g DW) | 6.90 Â± 2.61 c | 11.60 Â± 1.60 b | 19.32 Â± 1.14 a | 2.445e-12 *** |
|  | FRAP_Bound (µmol Fe2+ eq./g DW) | 42.56 Â± 13.58 b | 55.53 Â± 11.60 b | 137.18 Â± 12.01 a | 2.138e-14 *** |
| Total | TPC_Total (mg GAE/g DW) | 230.03 Â± 14.69 a | 209.96 Â± 10.97 b | 213.95 Â± 6.60 b | 0.002141 ** |
|  | TFC_Total (mg CAE/g DW) | 6.39 Â± 0.59 a | 6.17 Â± 1.27 a | 3.23 Â± 0.21 b | 1.542e-08 *** |
|  | DPPH_Total (mg Trolox eq./g DW) | 266.12 Â± 23.27 a | 275.73 Â± 19.64 a | 200.68 Â± 37.03 b | 9.024e-06 *** |
|  | FRAP_Total (µmol Fe2+ eq./g DW) | 1668.54 Â± 182.84 b | 1825.62 Â± 199.57 b | 2339.17 Â± 125.90 a | 4.35e-08 *** |

Statistical signiﬁcance is given as follows: * p < 0.05, ** p < 0.001, and *** p < 0.005, as compared to untreated controls. TPC, total phenolic content, mg GAE/ g DW; TFC, total flavonoid content, mg CAE /g DW; DPPH, 2,2-diphenyl-1-picrylhydrazyl, mg TE/ g DW; FRAP, ferric reducing antioxidant power, μmol Fe2+/g DW.

Table S2. The antioxidant properties (i.e., TPC, TFC, DPPH, and FRAP) for free, bound, and total phenolics of Kakadu extracts assisting with different extraction technologies.

|  | Variable | Microwave | Shaking | Ultrasound | p-value |
| --- | --- | --- | --- | --- | --- |
| Free | TPC_Free (mg GAE/g DW) | 209.11 Â± 10.40 a | 210.99 Â± 16.57 a | 215.58 Â± 15.77 a | 0.6286 n.s. |
|  | TFC_Free (mg CAE/g DW) | 4.86 Â± 1.70 a | 5.25 Â± 2.09 a | 4.48 Â± 1.82 a | 0.6879 n.s. |
|  | DPPH_Free (mg Trolox eq./g DW) | 227.47 Â± 53.38 a | 233.50 Â± 47.52 a | 243.74 Â± 43.05 a | 0.8339 n.s. |
|  | FRAP_Free (µmol Fe2+ eq./g DW) | 1916.48 Â± 336.91 a | 1781.46 Â± 312.39 a | 1900.12 Â± 260.04 a | 0.5976 n.s. |
| Bound | TPC_Bound (mg GAE/g DW) | 5.59 Â± 2.05 a | 6.16 Â± 2.15 a | 6.50 Â± 1.21 a | 0.4174 n.s. |
|  | TFC_Bound (mg CAE/g DW) | 0.40 Â± 0.15 a | 0.38 Â± 0.31 a | 0.42 Â± 0.18 a | 0.4295 n.s. |
|  | DPPH_Bound (mg Trolox eq./g DW) | 12.97 Â± 5.14 a | 11.62 Â± 6.40 a | 13.23 Â± 5.48 a | 0.8517 n.s. |
|  | FRAP_Bound (µmol Fe2+ eq./g DW) | 79.52 Â± 44.23 a | 74.48 Â± 46.93 a | 81.28 Â± 46.87 a | 0.8517 n.s. |
| Total | TPC_Total (mg GAE/g DW) | 214.70 Â± 9.43 a | 217.15 Â± 16.34 a | 222.08 Â± 15.69 a | 0.5395 n.s. |
|  | TFC_Total (mg CAE/g DW) | 5.25 Â± 1.61 a | 5.63 Â± 1.80 a | 4.90 Â± 1.69 a | 0.6648 n.s. |
|  | DPPH_Total (mg Trolox eq./g DW) | 240.43 Â± 48.74 a | 245.12 Â± 43.59 a | 256.97 Â± 40.16 a | 0.7757 n.s. |
|  | FRAP_Total (µmol Fe2+ eq./g DW) | 1996.00 Â± 369.01 a | 1855.94 Â± 350.76 a | 1981.40 Â± 305.58 a | 0.6404 n.s. |

Statistical signiﬁcance is given as follows: * p < 0.05, ** p < 0.001, and *** p < 0.005, as compared to untreated controls. TPC, total phenolic content, mg GAE/ g DW; TFC, total flavonoid content, mg CAE /g DW; DPPH, 2,2-diphenyl-1-picrylhydrazyl, mg TE/ g DW; FRAP, ferric reducing antioxidant power, μmol Fe2+/g DW.

Table S3. Predicted physicochemical, drug-likeness, ADMET and toxicokinetic properties of 24 compounds. (Please find in the excel document)

Each column is a compound; each row is a parameter. Values are RDKit-computed physicochemistry, ADMET-AI machine-learning predictions (classification endpoints reported as probabilities in [0, 1], regression endpoints in the stated units), and httk human toxicokinetics. Drug-likeness rules are reported as Yes (pass) / No (fail). For toxicity probabilities, lower values indicate lower predicted risk.

Parameter notes:

- Topological PSA (Å²): lower favours membrane permeability (≤140; ≤90 for CNS).

- LogP (RDKit): RDKit Crippen MolLogP; LogP (ADMET-AI) is the ML estimate.

- QED (drug-likeness): 0–1, higher is more drug-like.

- Synthetic accessibility (1-10): 1 = easy to synthesise, 10 = difficult.

- GI absorption (HIA): Yes if predicted HIA probability ≥ 0.5.

- Oral bioavailability (Ma): predicted probability of ≥20% oral bioavailability.

- Aqueous solubility (log mol/L): regressed log molar solubility (AqSolDB).

- Developability score (0-1): composite rank metric (see Methods); higher is better.

Abbreviations: LogP, octanol–water partition coefficient (lipophilicity); QED, quantitative estimate of drug-likeness; HIA, human intestinal absorption; BBB, blood–brain-barrier penetration; P-gp, P-glycoprotein; AMES, Ames mutagenicity assay; hERG, human ether-a-go-go-related gene (cardiac K+ channel) inhibition; DILI, drug-induced liver injury; PAINS, pan-assay interference compounds; prob, predicted probability (0-1).

Table S4. Target-compartment ADMET reachability screen of 26 compound-target pair(s) administered by the oral route (4 pharmacokinetically feasible; 4 carried forward to docking). (Please find in the excel document)

Each row is one compound-target pair. Feasibility is decided by a serial chain of barriers: a compound must (i) reach the target tissue (Axis A), (ii) reach the sub-cellular site, crossing the plasma membrane only when the target is intracellular (Axis B), and (iii) achieve adequate free (unbound) exposure. A pair is feasible only when all three are satisfied (axisA_pass AND axisB_pass AND exposure_pass).

Column groups:

- Localization: tissue_location (Axis A) and subcellular_class (Axis B) are derived from UniProt subcellular location, membrane topology and GO cellular-component annotations; membrane_crossing_required is True for intracellular/membrane targets.

- Key ADMET values: raw predictions used by the gates — admet_HIA_Hou (intestinal absorption), admet_Bioavailability_Ma, admet_Solubility_AqSolDB (log mol/L), admet_Caco2_Wang (permeability), admet_Pgp_Broccatelli (efflux), admet_BBB_Martins, pc_tpsa, pc_hbd, pc_mw, pc_mol_logp, admet_PPBR_AZ (plasma protein binding %), tk_fup_used (free fraction), admet_Half_Life_Obach, admet_VDss_Lombardo; plus safety flags admet_hERG/DILI/AMES/ClinTox.

- Per-parameter verdicts (*_pass): pass / fail / na (na = the gate did not apply for this pair's route or localization).

- failure_reasons: semicolon-separated identifiers of the gates that failed (or missing:<column> / admet_row_not_found / missing_structure).

Default thresholds (overridable): Axis A oral-systemic — HIA >= 0.5, Bioavailability_Ma >= 0.3, Solubility_AqSolDB >= -5 log mol/L, Pgp < 0.7; Axis A CNS — BBB >= 0.5, Pgp < 0.5, TPSA <= 90, HBD <= 3, MW <= 450; Axis B intracellular — Caco2 >= -5.5, logP 1.0-3.5, TPSA <= 120, HBD <= 5, MW <= 500, Pgp < 0.7 (organelle/mitochondrion additionally logP >= 2.0); membrane-embedded — logP >= 1.5; exposure — free fraction (fup) >= 0.001.

Abbreviations: ADMET, absorption-distribution-metabolism-excretion-toxicity; HIA, human intestinal absorption; BBB, blood-brain barrier; P-gp, P-glycoprotein; TPSA, topological polar surface area; HBD, hydrogen-bond donors; MW, molecular weight; logP, octanol-water partition coefficient; fup, fraction unbound in plasma; PPBR, plasma protein binding.

Table S5. Binding energy between selected phenolic compounds and core targets relevant to oxidative stress diseases including PTGS2, MMP2, TNF-alpha of human and canine.

| **Target** | **Animal model** | **Target uniprot** | **Ligand name** | **SMILES of ligand** | **Binding energy (kcal/mol)** |
| --- | --- | --- | --- | --- | --- |
| PTGS2 | Human | P35354 | 3,3',4'-Tri-O-methylellagic acid | COC1=C(C2=C3C(=C1)C(=O)OC4=C3C(=CC(=C4OC)OC)C(=O)O2)O | -4.848 |
| MMP2 | Human | P08253 | Arjunolic acid | C[C@@]12CC[C@@H]3[C@@]([C@H]1CC=C4[C@]2(CC[C@@]5([C@H]4CC(CC5)(C)C)C(=O)O)C)(C[C@H]([C@@H]([C@@]3(C)CO)O)O)C | -6.622 |
| MMP2 | Human | P08253 | Linolenic acid | CC/C=C\C/C=C\C/C=C\CCCCCCCC(=O)O | -7.059 |
| TNF-alpha | Human | P01375 | Linolenic acid | CC/C=C\C/C=C\C/C=C\CCCCCCCC(=O)O | -4.349 |
| PTGS2 | Dog | A0A8I3MVI5 | 3,3',4'-Tri-O-methylellagic acid | COC1=C(C2=C3C(=C1)C(=O)OC4=C3C(=CC(=C4OC)OC)C(=O)O2)O | -8.641 |
| MMP2 | Dog | A0A8I3MGQ5 | Arjunolic acid | C[C@@]12CC[C@@H]3[C@@]([C@H]1CC=C4[C@]2(CC[C@@]5([C@H]4CC(CC5)(C)C)C(=O)O)C)(C[C@H]([C@@H]([C@@]3(C)CO)O)O)C | -7.667 |
| MMP2 | Dog | A0A8I3MGQ5 | Linolenic acid | CC/C=C\C/C=C\C/C=C\CCCCCCCC(=O)O | -4.985 |
| TNF-alpha | Dog | P51742 | Linolenic acid | CC/C=C\C/C=C\C/C=C\CCCCCCCC(=O)O | -3.976 |
