## Supplementary figures for "Optimized Kakadu Plum Extracts Inhibit Intracellular Oxidative Stress in Canine Small Intestinal Cell Model"

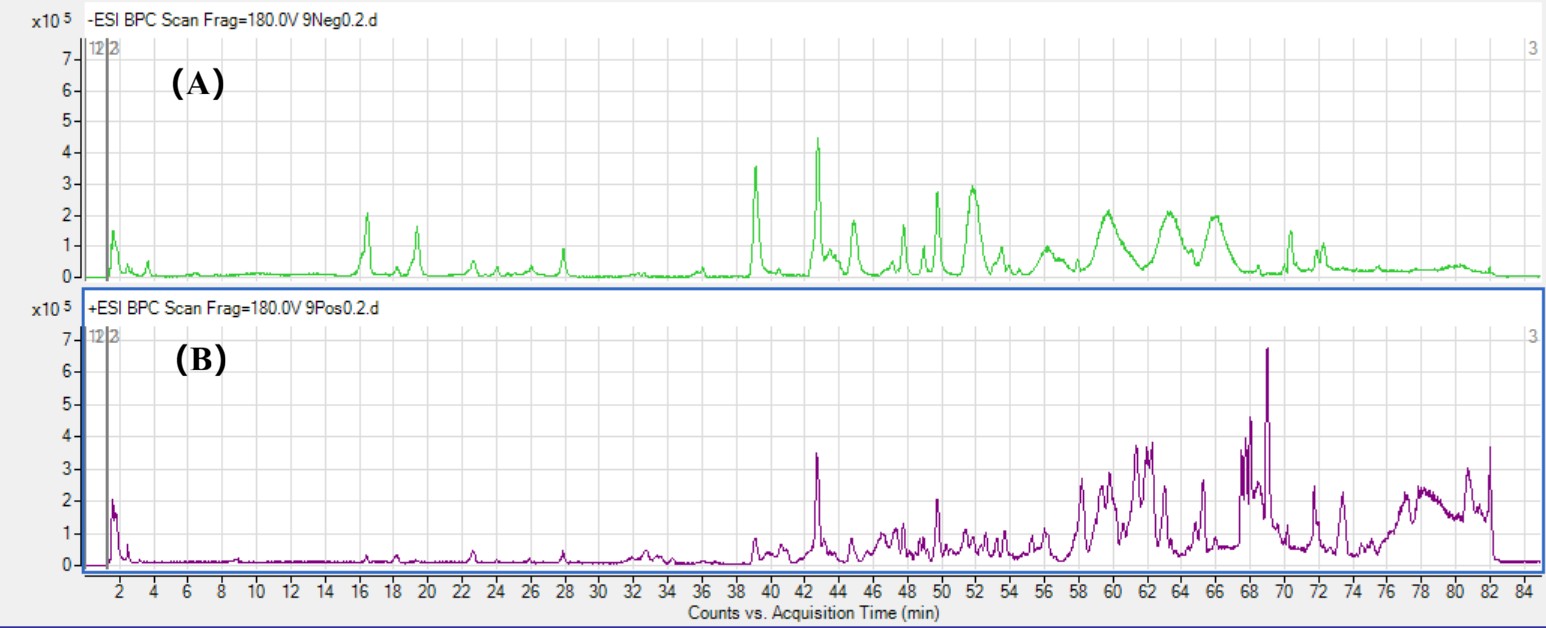

Figure S1. HPLC-ESI-QTOF-MS/MS Spectra of Kakadu plum ethanol extract. (A): Negative mode. (B): Positive mode.

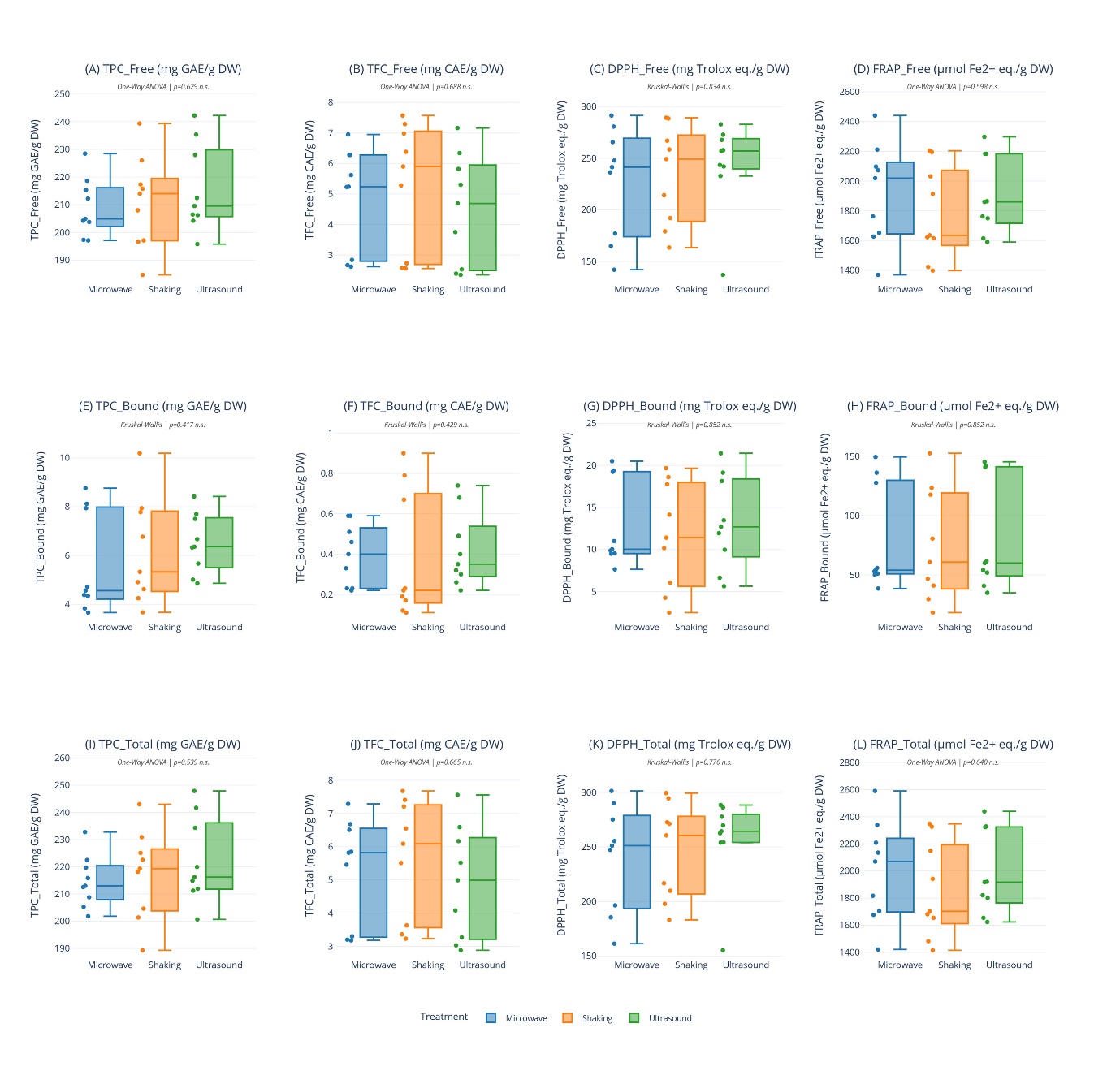

Figure S2. The antioxidant properties (i.e., TPC, TFC, DPPH, and FRAP) for free, bound, and total phenolics of Kakadu extracts assisting with different extraction technologies. A: TPC of free phenolics of Kakadu extracts. B: TFC of free phenolics of Kakadu extracts. C: DPPH of free phenolics of Kakadu extracts. D: FRAP of free phenolics of Kakadu extracts. E: TPC of bound phenolics of Kakadu extracts. F: TFC of bound phenolics of Kakadu extracts. G: DPPH of bound phenolics of Kakadu extracts. H: FRAP of bound phenolics of Kakadu extracts. I: TPC of total phenolics of Kakadu extracts. J: TFC of total phenolics of Kakadu extracts. K: DPPH of total phenolics of Kakadu extracts. L: FRAP of total phenolics of Kakadu extracts.TPC, total phenolic content, mg GAE/ g DW; TFC, total flavonoid content, mg CAE /g DW; DPPH, 2,2-diphenyl-1-picrylhydrazyl, mg TE/ g DW; FRAP, ferric reducing antioxidant power, μmol Fe2+/g DW.

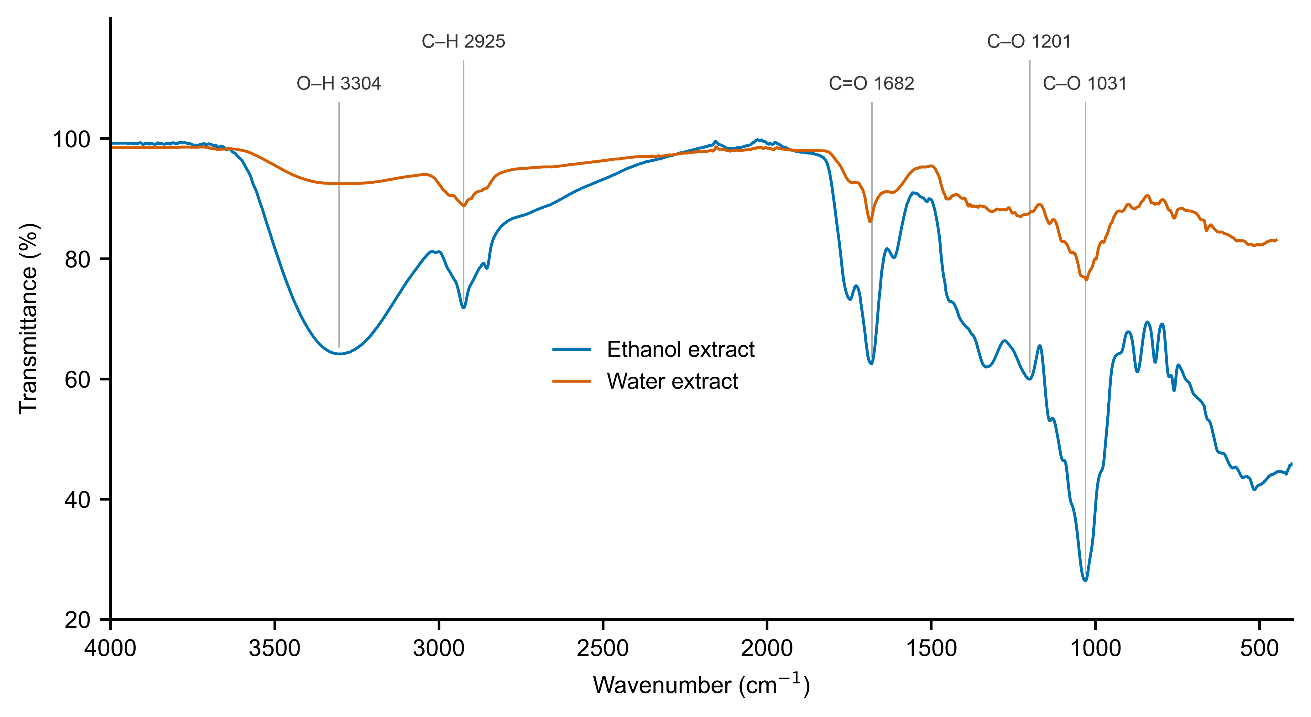

Figure S3. FTIR spectra of water and ethanol extract of Kakadu plum.

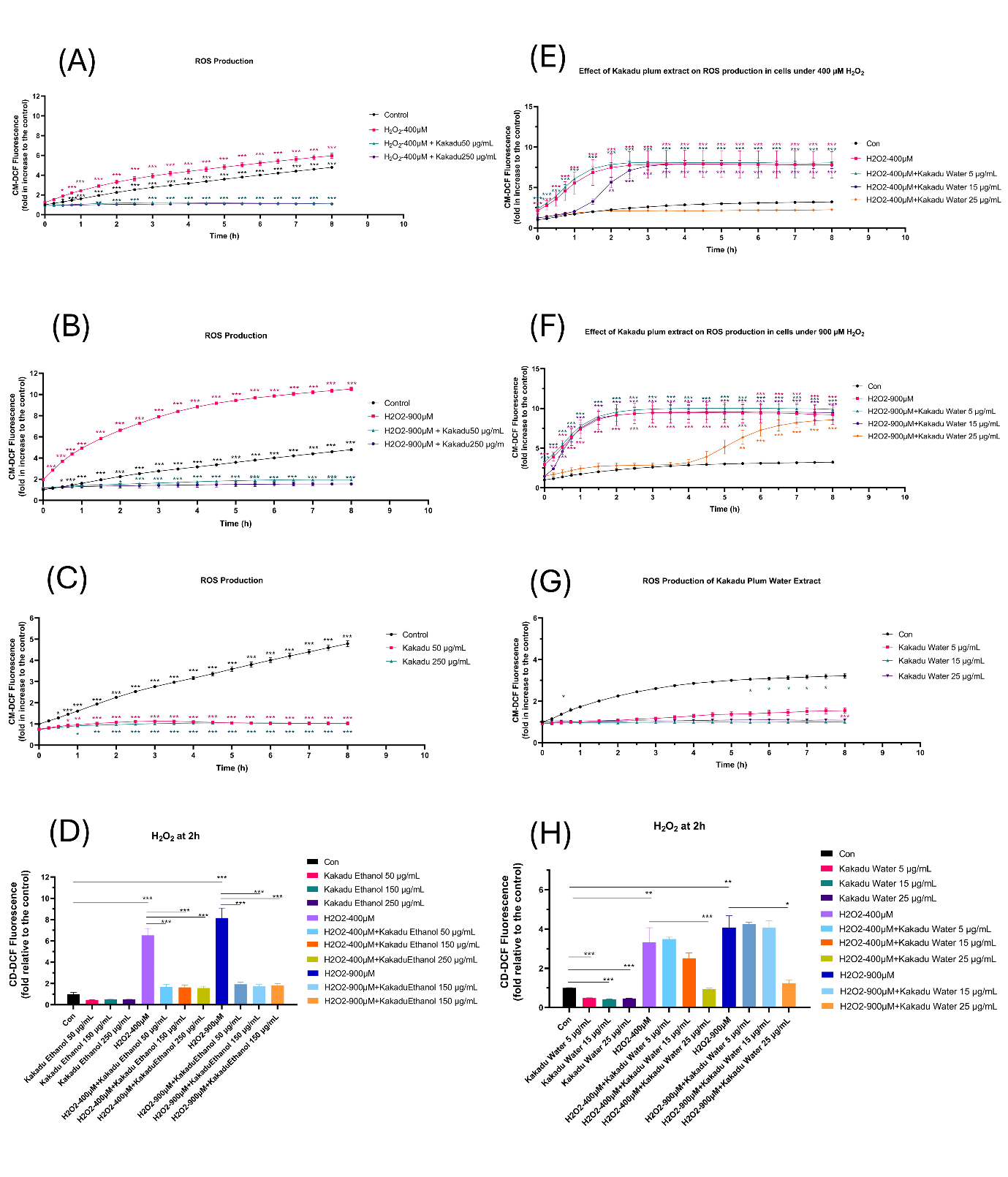

Figure S4. Intracellular ROS production of cells under treatment of water and ethanol extract of Kakadu plum.

A: The effect of water extract of Kakadu plum on the ROS production under mild (400 µM H₂O₂) oxidative stress model. B: The effect of water extract of Kakadu plum on the ROS production under severe (900 µM H₂O₂) oxidative stress model. C: The effect of water extract of Kakadu plum on the ROS production under no oxidative stress. D: The inhibitory effect of water extract of Kakadu plum on the ROS production under oxidative stress for 2h. E: The effect of ethanol extract of Kakadu plum on the ROS production under mild (400 µM H₂O₂) oxidative stress model. F: The effect of ethanol extract of Kakadu plum on the ROS production under severe (900 µM H₂O₂) oxidative stress model. G: The effect of ethanol extract of Kakadu plum on the ROS production under no oxidative stress. H: The inhibitory effect of ethanol extract of Kakadu plum on the ROS production under oxidative stress for 2h.

Cells were seeded at densities of 2.0×104 in 96-well plates and incubated for 24 h with increasing concentrations of H₂O₂. The data are normalized as the relative response of treated cells as compared to the untreated control from triplicate experiments. Statistical signiﬁcance is given as follows: * p < 0.05, ** p < 0.001, and *** p < 0.005, as compared to untreated controls.
